# Age and spatio-temporal variations in food resources modulate stress-immunity relationships in three populations of wild roe deer

**DOI:** 10.1101/2022.02.15.480486

**Authors:** Jeffrey Carbillet, Marine Hollain, Benjamin Rey, Rupert Palme, Maryline Pellerin, Corinne Regis, Anne Geffré, Jeanne Duhayer, Sylvia Pardonnet, François Debias, Joël Merlet, Jean-François Lemaître, Hélène Verheyden, Emmanuelle Gilot-Fromont

**Affiliations:** Université de Toulouse, INRAE, CEFS, Castanet Tolosan, 31326, France; LTSER ZA PYRénées GARonne, Auzeville-Tolosane, 31320, France; Université de Lyon, VetAgro Sup, Marcy-l’Etoile, 69280, France; Institute of Ecology and Earth Sciences, University of Tartu, Tartu, 51014, Estonia; Université de Lyon, Université Lyon 1, UMR CNRS 5558, Villeurbanne Cedex, 69100, France; Office Français de la biodiversité, Direction de la Recherche et de l’Expertise, Unité Ongulés Sauvages, Gières, 38610, France; Unit of Physiology, Pathophysiology, and Experimental Endocrinology, Department of Biomedical Sciences, University of Veterinary Medicine, Vienna, 1210, Austria; Equipe de Biologie médicale-Histologie, CREFRE, Inserm-UPS-ENVT, Toulouse, 31000, France

**Keywords:** stress hormones, ecophysiology, innate immunity, adaptive immunity, faecal glucocorticoid metabolites, trade-off

## Abstract

Living in variable and unpredictable environments, organisms face recurrent stressful situations. The endocrine stress response, which includes the secretion of glucocorticoids, helps organisms to cope with these perturbations. Although short-term elevations of glucocorticoid levels are often associated with immediate beneficial consequences for individuals, long-term glucocorticoid elevation can compromise key physiological functions such as immunity. While laboratory works highlighted the immunosuppressive effect of long-term elevated glucocorticoids, it remains largely unknown, especially in wild animals, whether this relationship is modulated by individual and environmental characteristics. In this study, we explored the co-variation between baseline cortisol levels, assessed non-invasively using faecal cortisol metabolites (FCMs), and 12 constitutive indices of innate, inflammatory, and adaptive immune functions, in wild roe deer living in three populations with contrasting environmental conditions. Using longitudinal data on 564 individuals, we further investigated whether age and spatio-temporal variations in the quantity and quality of food resources affect the relationship between FCMs and immunity. Negative covariation with glucocorticoids was evident only for innate and inflammatory markers of immunity, while adaptive immunity appeared to be positively or not linked to glucocorticoids. In addition, the negative covariations were generally exacerbated, or revealed, in individuals facing harsh environmental constraints and in old individuals. Therefore, our results highlight the importance of measuring multiple immune markers of immunity in individuals from contrasted environments to unravel the complex relationships between glucocorticoids and immunity in wild animals. Our results also help explain conflicting results found in the literature and could improve our understanding of the long-term consequences of elevated glucocorticoid levels on disease spread and population dynamics.

## 1. Introduction

The neuroendocrine stress response helps animals to cope with recurrent stressful situations in natural environments (Sapolsky et al., 2000; Wingfield and Romero, 2001). Exposure to stressors stimulates the hypothalamic-pituitary-adrenal (HPA) axis and leads, among others, to an increase in the secretion of glucocorticoids by the adrenocortex (Sapolsky et al., 2000; Sheriff et al., 2011). Glucocorticoids, which contribute to the control of an individual’s energy balance through acquisition, storage and mobilization, also coordinate the body’s overall response to stressors through metabolic changes that depend on their concentration and duration of secretion. (Hau et al., 2016). This hormonal cascade is then subjected to negative feedback from glucocorticoids on their own secretion (Romero, 2004; Sheriff et al., 2011), facilitating a return to a baseline level that ensures maintenance of daily activities according to the current life-history stage (Möstl and Palme, 2002; Wingfield and Sapolsky, 2003). While short-term elevation of glucocorticoid levels promotes survival (Sapolsky et al., 2000; Breuner et al., 2008), chronic elevation of glucocorticoids may alter physiological functions and ultimately compromise both survival and reproductive success on the long-run (Boonstra, 2005). Among these functions, the potential immunosuppressive action of glucocorticoids is one of the most discussed effects of chronic stress (Martin, 2009).

Immunity is a key physiological function of vertebrates: it includes innate and adaptive components (Stanley, 2002), each comprising cellular and humoral effectors (Stanley, 2002). While innate immunity sets up rapidly (within hours) and is mostly non-specific, adaptive (memory-based) immunity deals with repeated infections and selectively eliminates pathogens (Lee, 2006). Like any other physiological function, the maintenance and functioning of the immune system requires energy (Lee, 2006; Martin, 2009). Hence, immunity has been hypothesised to trade-off against other energy demanding physiological functions (Martin, 2009). Glucocorticoids may mediate these trade-offs, with elevation of these hormones redirecting the energy away from immunity towards functions promoting immediate survival but with deleterious effects in the long term (Lee, 2006; Martin, 2009). However, costs vary depending on the stage (development, maintenance, or activation) and component of immunity (Klasing, 2004, Lee, 2006). The adaptive immune response is thought to have a lower activation energy cost than the innate immune response, the inflammatory response entailed by innate response being particularly energy demanding (McDade, 2016). Therefore, differential effects of trade-off mediated by glucocorticoids can be expected between the components of immunity.

So far, most empirical studies documenting a detrimental effect of chronic stress on immune functions have been conducted on laboratory or domestic animals (e.g. Dhabhar et al., 1994; Dhabhar et al., 1995; Wada et al., 2010; Moazzam et al., 2012) and much less studies have been conducted in wild populations (e.g. Bourgeon and Raclot, 2006; Brooks and Mateo, 2013; Josserand et al., 2020). In wild populations, exposure to fluctuating environmental conditions is more intense than in captivity in terms of resources, variation of temperature, predator risk or disease threats for instance, which could lead to different outcomes regarding the relationship between glucocorticoids and immunity. In addition, a large body of literature shows that diverse internal and external factors could influence both immunity (Lee, 2006; Martin, 2009) and glucocorticoid secretion (Hau et al., 2016). For instance, poor environments (in terms of resource quantity or quality) can lead to both increased glucocorticoid levels (Fokidis et al., 2012; Carbillet et al., 2020) and decreased concentrations of immune parameters (Dhabhar et al., 1994; Dhabhar et al., 1995), without any causal relationships. It has also been shown that baseline glucocorticoid levels are related to age (Sapolsky et al., 1983) as are several immune parameters (e.g. immunosenesence, see Nussey et al., 2012; Cheynel et al., 2017). To date, the contribution of these individual and environmental factors has not been investigated and their role in modulating the relationship between immunity and glucocorticoids remains poorly understood.

The aim of this study was to analyse whether the relationship between glucocorticoid levels and immunity is affected by both individual and environmental factors in three wild populations of roe deer (*Capreolus capreolus*) living in habitats with contrasting environmental conditions. Baseline glucocorticoid levels were assessed non-invasively by measuring faecal cortisol metabolites (FCMs), which reflect overall circulating glucocorticoid levels that an individual has experienced over a particular time-period which is species-specific (Palme et al., 2005; Sheriff et al., 2011; Palme, 2019). The immune function was assessed the same years than FCMs by measuring twelve immune parameters encompassing the innate (neutrophils, eosinophils, basophils, monocytes, hemagglutination, hemolysis), inflammatory (alpha-1-globulin, alpha-2-globulin, beta-globulin, haptoglobin) and adaptive (lymphocytes, gamma-globulin) immunity (Cheynel et al. 2017).

Based on current knowledge, we expected that individuals with higher baseline glucocorticoid levels would exhibit weaker overall immunity than those with lower baseline glucocorticoid levels. More precisely, we expected (prediction 1 that high levels of FCMs would be associated with low levels of parameters measuring innate (cellular and humoral) immunity (neutrophils, eosinophils, basophils, monocytes, hemagglutination, hemolysis), as well as inflammation (alpha-1-globulin, alpha-2-globulin, beta-globulin and haptoglobin) due to their high activation cost (Dhabhar et al., 2012; Brooks and Mateo, 2013; McDade et al., 2016). On the opposite, we expected a weaker relationship between FCMs and cellular and humoral adaptive immune parameters (lymphocyte and gamma-globulin concentrations) due to their lower activation cost (Klasing, 2004, Lee, 2006). In addition, we expected (prediction 2) that the negative relationship between FCMs and immune parameters would appear, or be exacerbated, in individuals facing poor environmental conditions such as low availability in food resources, compared to individuals living in favourable environments (Hau et al., 2016). Finally, we expected (prediction 3) that the negative relationship between FCM levels and immune parameters would appear, or be exacerbated, in older individuals compared with younger ones, due to impairment of both the adrenocortical stress response (Sapolsky et al., 1983) and the immune response in old individuals (Cheynel et al., 2017).

## 2. Material and methods

### 2.1 Study populations

This study was conducted on three wild populations of roe deer living in contrasting habitats. First, the Trois-Fontaines population is located in an enclosed forest (1,360 hectares) in the north-east of France (48°43’ N, 4°55’ E). The climate is continental, and the soil is particularly rich, making it a very productive forest offering a homogeneous and high-quality resources habitat for roe deer (Pettorelli et al., 2006). Second, the Chizé population is located in an enclosed forest (2,614 hectares) in western France (46°50’ N, 0°25’ W). The climate is temperate oceanic, with Mediterranean influences. Due to poor soil quality and frequent summer droughts, the forest productivity is low compared to Trois-Fontaines (Pettorelli et al., 2006), making it a relatively poor-quality habitat for roe deer in terms of resources (Gaillard et al., 1993). At a finer scale, three sectors are distinguished in this population according to the quantity and quality of resources (Pettorelli et al., 2001). Sector 1 is composed of oaks (*Quercus spp*.) and hornbeams (*Carpinus betulus*) and is considered to be of better quality than the other two sectors, sector 2 is composed of oaks and Montpellier maples (*Acer monspessulanum*) and is considered to be of intermediate quality, and sector 3, composed of beeches (*Fagus sylvatica*) is the sector of worse quality. Pettorelli and colleagues (2003) reported that these differences in habitat composition result in differences in roe deer body mass, with juveniles in sector 1 being on average 2 kg heavier than those in sector 3. Finally, the Aurignac population is located in an agricultural landscape (10,000 hectares), in south-western France (43°13’ N, 0°52’ E) and is part of a Long-Term Socio-Ecological Research platform called ZA PYGAR. This site is exposed to an oceanic climate, with summer droughts (Hewison et al., 2007). It provides a highly heterogeneous environment with a fragmented landscape composed of forests, grassland and cultivated fields (see Martin et al., 2018 for details).

This study site provides overall high-quality habitat for roe deer (as Trois-Fontaines) but can also be divided into three sectors according to resource quality and habitat openness (see Morellet et al., 2009 for details). The most open habitats (sector 1) offer more important and high-quality food resources for roe deer during most of the year (Abbas et al., 2011), but can also be a source of higher exposure to stressors (Bonnot et al., 2013), such as road and human dwellings leading to higher FCM levels (Carbillet et al., 2020), compared to the partially wooded area (sector 2) and woodland (sector 3). Hewison and colleagues (2009) have shown a beneficial effect of landscape openness on roe deer reproduction, demographic performance, and juveniles body mass, with juveniles in sector 3 weighing on average 2.0 kg less than in sector 2, and 3.1 kg less than in sector 1 (Hewison et al., 2009).

### 2.2 Data collection

As part of long-term capture-mark-recapture programs initiated in 1975, 1977 and 1996, in Trois-Fontaines, Chizé and Aurignac respectively, 6 to 12 days of drive net captures are organised between December and March each year. At each capture session, between 30 and 100 beaters push roe deer towards nets deployed over 4 km. Once captured, each animal is marked, weighed (to the nearest 100 g), sexed, and age is determined by tooth eruption patterns in Aurignac (Hewison et al., 1999), with 2 age classes: juveniles (< 1 year) and adults (> 1 year). For the Trois-Fontaines and Chizé populations, the exact age (in years) is known since the individuals were all captured and marked during their first year of life when the age can be estimated without error. Blood samples are taken from the jugular vein (up to 1 mL/kg) since 2009 in Aurignac and since 2010 in Trois-Fontaines and Chizé. Part of the collected blood is transferred to a dry tube, centrifuged, and the serum conserved at -20°C for biochemical analyses. The remaining blood is transferred to a tube containing ethylenediaminetetraacetic acid (EDTA) for further determination of immune parameters (see below). For each individual, faeces are collected rectally and stored at -20°C until extraction.

### 2.3 Measurement of FCMs

Faecal cortisol metabolites were extracted following the protocol developed by Palme and colleagues (2013). Briefly, for each individual, 0.5 g of faeces were suspended in 5 mL of 80% methanol, vortexed for 30 minutes and centrifuged for 15 min at 2500 g. The supernatants were diluted 1:10 with assay buffer before FCM concentrations were determined with a group-specific 11-oxoaetiocholanolone enzyme immunoassay as previously described (Möstl et al., 2002) and validated for roe deer (Zbyryt et al., 2017). Measurements were carried out in duplicate. Intra- and inter-assay coefficients of high and low concentration pool samples were less than 10% and 15%, respectively. Results are expressed as nanograms per gram of wet faeces (ng/g).

### 2.4 Measurement of immune parameters

#### 2.4.1 Innate cellular immunity

White blood cells (WBC) count was carried out by impedance technology (ABC Vet automaton, Horiba Medical) and the proportions of each WBC type (neutrophils, monocytes, lymphocytes, eosinophils and basophils) were quantified under microscope (x1000) by counting the first 100 WBC on blood smears, previously stained with a May-Grünwald and Giemsa solution (see Houwen, 2001 for more details). The concentrations of each type of leukocyte were then determined as (WBC*parameter cells count/100).

#### 2.4.2 Innate humoral immunity

Innate humoral immunity was assessed by measuring circulating levels of natural antibodies (NAbs) and complement activity of serum. The concentration of NAbs was measured by the hemagglutination test (HA) that measures NAbs ability to agglutinate exogenous cells (Matson et al., 2005). The complement is a group of proteins that acts in a chain reaction and causes lysis of exogenous cells in presence of an antigen-antibody complex. It is revealed by the ability of proteins to induce hemolysis (HL) (Matson et al., 2005). The HA/HL protocol developed by Matson et al., (2005) for birds has been previously adapted for roe deer (Gilot-Fromont et al., 2012).

#### 2.4.3 inflammatory markers

Inflammatory status was evaluated using levels of alpha-1, alpha-2, and beta globulins. The total concentrations of these proteins (in g/L) were quantified by a refractometer followed by an electrophoresis on agarose gel, using an automaton (HYDRASYS). Haptoglobin concentration (in mg/mL), which belongs to the alpha-2-globulin fraction and reflect infection or chronic inflammation, was also measured. Analyses were performed with a Konelab 30i PLC (Fisher Thermo Scientific) which operates on the principle of spectrophotometry. Unlike other immune parameters, haptoglobin was measured only on the Trois-Fontaines and Chizé samples.

#### 2.4.4 Adaptive cellular immunity

Adaptive cellular immunity was assessed using lymphocyte concentration, determined by the leukocyte count described above, which includes both B and T lymphocytes.

#### 2.4.5 Adaptive humoral immunity

The adaptive humoral immunity component was assessed using concentration of gamma globulins, obtained by the protein analysis described above for other globulins. Gamma globulin concentration has been used as an estimator of total antibodies, since they are essentially composed of circulating antibodies (Stockham and Scott, 2008).

### 2.5 Statistical analyses

Due to conservation issues of faeces, the available data encompassed different year ranges according to the population. Data were available for the years 2010 to 2019 in Trois-Fontaines, while only the years 2013, 2014 and 2016 to 2019 were available in Chizé. For the Aurignac population, neutrophils, esosinophils, basophils, monocytes and lymphocytes concentrations were available for the years 2012 to 2017. For hemagglutination and hemolysis data were available from the years 2013 to 2017, while for gamma-, alpha-1-, alpha-2- and beta-globulin, data were available for the years 2014 to 2017. Consequently, the number of observations differed according to the population and immune marker considered. In Aurignac, analyses were carried out on 144 to 188 observations, while they were conducted on 325 to 414 in Chizé, and 276 to 303 in Trois-Fontaines. Overall, we implemented 35 series of models, including 12 series for the populations of Trois-Fontaines and Chizé and 11 series for Aurignac (due to the absence of haptoglobin assays).

For each of the 12 immune parameters, we ran separate analyses for each of the three populations, using linear mixed effect models (LMMs). Each immune trait was taken as a response variable and some of them (concentrations of eosinophils, basophils, monocytes, lymphocytes, haptoglobin, alpha-1, alpha-2 and beta globulins) were transformed as log(x+1) to ensure normality of model residuals. Model selection was carried out by adjusting a reference model (see below) that included all biologically relevant variables and interactions to test our hypotheses. We then compared this model with all its sub-models. Each continuous explanatory variable (except for age) was centred by population to obtain estimates corresponding to average values of the parameters in the corresponding population. The reference model included:

i. Individual variables: baseline cortisol level (FCMs), log-transformed and centred around the mean (mean value of the variable that is subtracted from every value), sex, body weight (in kg, centred), and age. In Aurignac, age was considered with two classes (juveniles and adults). In Chizé and Trois-Fontaines, age in years was considered to have either a linear effect, a quadratic function, or a threshold effect) based on a previous study carried out on these two populations (Cheynel et al., 2017).
ii. Environmental variables: in Chizé and Aurignac, we considered the sector of capture as a marker of local resources quality and quantity. To account for temporal variations of resources among years, we used a centred year quality index. The quality of a given year was indexed using the average weight (in kg, centred) of juveniles caught the following winter (Pettorelli et al., 2003).
iii. Methodological variables: the time between capture and blood sampling is known to influence the level of certain immune parameters such as neutrophils and lymphocytes (Carbillet et al., 2019) and was thus taken into account (thereafter, delay, in minutes, centred). In addition, we included the Julian date of capture in our models to control for potential among-individual differences in immune parameters, body weight and FCMs due to the timing of sampling.

Several plausible interactions based on our hypotheses were also included in our reference model. First, the interaction between FCMs and age, to investigate a possible modification of the relationship with advancing age (prediction 3). Second, we considered interactions between FCMs and sector, and between FCMs and quality of the year, to account for a potential modulation of the relationship under poorer environments or years of poorer quality (prediction 2). The interaction between FCMs and sex was also included to control for a possible modification of the relationship according to sex, as females generally allocate much more to immunity than males in the wild (Metcalf et al., 2019). We also included an interaction between FCMs and body weight to control for a possible modification of the relationship in individuals with the poorest physical condition (Hau et al., 2016). Finally, an interaction between sex and age was also included in neutrophil models for the Trois-Fontaines and Chizé populations because a previous study on roe deer from these populations showed that neutrophil profile was affected by age in a sex-specific manner (Cheynel et al., 2017). This interaction was also included in the Aurignac population for all immune parameters, as no previous data was available on the sex-specific effect of age on other parameters in this population. Individual’s identity and year of capture were included as random effects to avoid pseudo-replication problems (Hurlbert, 1984) and to control for unexplained variance due to among-individual differences and inter-annual variation. For the Chizé and Trois-Fontaines populations, the birth cohort is known and was also included as a random effect to account for potential differences between individuals in the environmental conditions they experienced early in life, which may have persistent effects on their phenotype (Douhard et al., 2014).

As a result, our (general) reference models read as follow:

Immune parameter = f(FCMs + year quality + sector + sex + age + weight + delay + julian date + FCMs*sector + FCMs*year quality + FCMs*sex + FCMs*age (either linear, quadratic or threshold) + FCMs*weight + 1|individual + 1|year of capture + 1|birth cohort).

To select the best models describing variation of each immune parameter, each reference model was compared to all its sub-models (N=358 models overall) using a model selection method based on the second-order Akaike’s information criteria (AICc, Burnham and Anderson, 2003). Models with a difference in AICc (ΔAICc) > 2 units from the best model were considered to have less support, following Burnham and Anderson (2003). In addition, we removed models within two AICc units of the top model that differed from a higher-ranking model by the addition of one or more parameters. These were rejected as uninformative, as recommended by Arnold (2010) and Richards (2008). Using the selected models, we then applied a conditional model averaging procedure to estimate parameters. We calculated AICc weights (AICcw) to measure the relative likelihood that a given model was the best among the set of fitted models and goodness-of-fit was assessed by calculating marginal (R2m) and conditional (R2c) variance using the r.squaredGLMM function of the MuMIn package (Barton, 2016). When interactions between FCMs and age, year quality, sectors, body weight, or sex were significant, we used F-tests in order to test prediction 1 of a relationship between FCMs and immunity. For each parameter in each population, the normality of residuals was tested (Shapiro-Wilk test) and visually assessed. All analyses were performed using R version 3.5.1 (R Development Core Team 2018) and using the lmer function (lme4 package, Bates et al., 2014) and the dredge function (MuMIn package, Barton, 2016).

## 3. Results

The text below describes the covariations between FCMs and immune parameters, and their variations according to the availability and quantity of food resources, age, sex, and body weight of roe deer (Table 1, 2 & 3). The full results of the model selection procedure are given in Tables S1, S2 and S3. A summary of these results is provided in Table S4. Correlation matrix between each immune parameters for each population are provided in Table S5.

**Table 1.** Parameters of linear mixed-effect models selected for each immune parameters of the Aurignac population. R2m and R2c correspond respectively to the marginal and conditional variance explained by the model, CI corresponds to the upper and lower limits of the 95% confidence interval, and n represents the number of observations per analysis. See the material and methods section for a full definition of model sets and explanation regarding the difference in the number of observations.

| Immune trait | Parameter | Estimate | CI |
| --- | --- | --- | --- |
| **Innate immunity** |  |  |  |
| **Neutrophils** (n = 188)<br>R<sup>2</sup><sub>m</sub> = 0.07 ; R<sup>2</sup><sub>c</sub> = 0.71 | Intercept | 6.07 | 5.75 to 6.39 |
|  | Delay | 0.006 | 0.002 to 0.01 |
|  | FCMs | -0.51 | -0.94 to -0.08 |
| **Eosinophils** (n = 185)<br>R<sup>2</sup><sub>m</sub> = 0.12 ; R<sup>2</sup><sub>c</sub> = 0.46 | Intercept | 0.05 | 0.04 to 0.07 |
|  | Age (juvenile) | -0.03 | -0.06 to -0.0001 |
|  | Julian date | 0.001 | 0.0003 to 0.002 |
|  | Delay | -0.0002 | -0.0003 to -0.00002 |
|  | Weight | -0.005 | -0.009 to -0.0008 |
|  | Year quality | -0.05 | -0.08 to -0.01 |
|  | Sex (male) | -0.02 | -0.04 to 0.003 |
| **Basophils** (n = 188)<br>R<sup>2</sup><sub>m</sub> = 0.04 ; R<sup>2</sup><sub>c</sub> = 0.32 | Intercept | 0.05 | 0.03 to 0.07 |
|  | Delay | -0.0001 | -0.003 to 0.00001 |
|  | Sector (2) | 0.01 | -0.01 to 0.04 |
|  | Sector (3) | 0.03 | 0.003 to 0.06 |
| **Monocytes** (n = 187)<br>R<sup>2</sup><sub>m</sub> = 0.01 ; R<sup>2</sup><sub>c</sub> = 0.01 | Intercept | 0.11 | 0.10 to 0.12 |
|  | Year quality | -0.04 | -0.09 to 0.007 |
| **Hemagglutination** (n = 168)<br>R<sup>2</sup><sub>m</sub> = 0.12 ; R<sup>2</sup><sub>c</sub> = 0.74 | Intercept | 3.64 | 3.15 to 4.16 |
|  | FCMs | -0.15 | -0.37 to 0.08 |
|  | Year quality | -1.54 | -3.49 to 0.39 |
|  | FCMs*Year quality | 0.96 | 0.07 to 1.86 |
| **Hemolysis** (n = 170)<br>R<sup>2</sup><sub>m</sub> = 0.31 ; R<sup>2</sup><sub>c</sub> = 0.66 | Intercept | 2.89 | 2.20 to 3.58 |
|  | FCMs | -0.19 | -0.43 to 0.05 |
|  | Year Quality | -3.12 | -5.84 to -0.41 |
|  | **Inflammatory markers** |  |  |
| **Alpha-1 globulins** (n = 146)<br>R<sup>2</sup><sub>m</sub> = 0.35 ; R<sup>2</sup><sub>c</sub> = 0.38 | Intercept | 1.40 | 1.36 to 1.44 |
|  | Julian date | 0.002 | 0.0003 to 0.003 |
|  | FCMs | 0.03 | -0.01 to 0.07 |
|  | Sector (2) | -0.03 | -0.08 to 0.01 |
|  | Sector (3) | -0.07 | -0.12 to -0.02 |
|  | Weight | -0.02 | -0.03 to -0.01 |
|  | Sex (Male) | 0.07 | 0.03 to 0.11 |
|  | FCMs*Sector (2) | 0.06 | -0.01 to 0.13 |
|  | FCMs*Sector (3) | -0.03 | -0.11 to 0.04 |
|  | FCMs*Weight | -0.006 | -0.01 to 0.002 |
|  | FCMs*Sex (male) | -0.05 | -0.11 to 0.01 |
| **Alpha-2 globulins** (n = 144)<br>R<sup>2</sup><sub>m</sub> = 0.35 ; R<sup>2</sup><sub>c</sub> = 0.38 | Intercept | 1.67 | 1.64 to 1.70 |
|  | Julian date | 0.0008 | -0.0002 to 0.002 |
| **Beta globulins** (n = 147)<br>R<sup>2</sup><sub>m</sub> = 0.10 ; R<sup>2</sup><sub>c</sub> = 0.13 | Intercept | 1.95 | 1.91 to 1.99 |
|  | Age (juvenile) | -0.16 | -0.25 to -0.07 |
|  | FCMs | 0.03 | -0.009 to 0.08 |
|  | Weight | -0.02 | -0.03 to -0.004 |
|  | **Adaptive immunity** |  |  |
| **Gamma globulins** (n = 147)<br>R<sup>2</sup><sub>m</sub> = 0.24 ; R<sup>2</sup><sub>c</sub> = 0.72 | Intercept | 13.16 | 12.05 to 14.28 |
|  | Delay | -0.006 | -0.01 to 0.0003 |
|  | Age (juvenile) | -3.47 | -5.12 to -1.81 |
|  | FCMs | 1.58 | 0.40 to 2.76 |
|  | Sector (2) | -1.13 | -2.31 to 0.05 |
|  | Sector (3) | -0.52 | -1.88 to 0.85 |
|  | Weight | -0.33 | -0.56 to -0.10 |
|  | Year quality | 1.75 | 0.20 to 3.30 |
|  | Sex (male) | 0.47 | -0.62 to 1.55 |
|  | FCMs*Sector (2) | -2.68 | -4.38 to -0.98 |
|  | FCMs*Sector (3) | -2.11 | -3.87 to -0.35 |
|  | FCMs*Sex (male) | -2.10 | -3.62 to -0.57 |
| **Lymphocytes** (n = 188)<br>R<sup>2</sup><sub>m</sub> = 0.24 ; R<sup>2</sup><sub>c</sub> = 0.72 | Intercept | 1.28 | 1.21 to 1.35 |
|  | Weight | -0.02 | -0.03 to -0.01 |
|  | Sex | -0.07 | -0.14 to -0.002 |

**Table 2.**
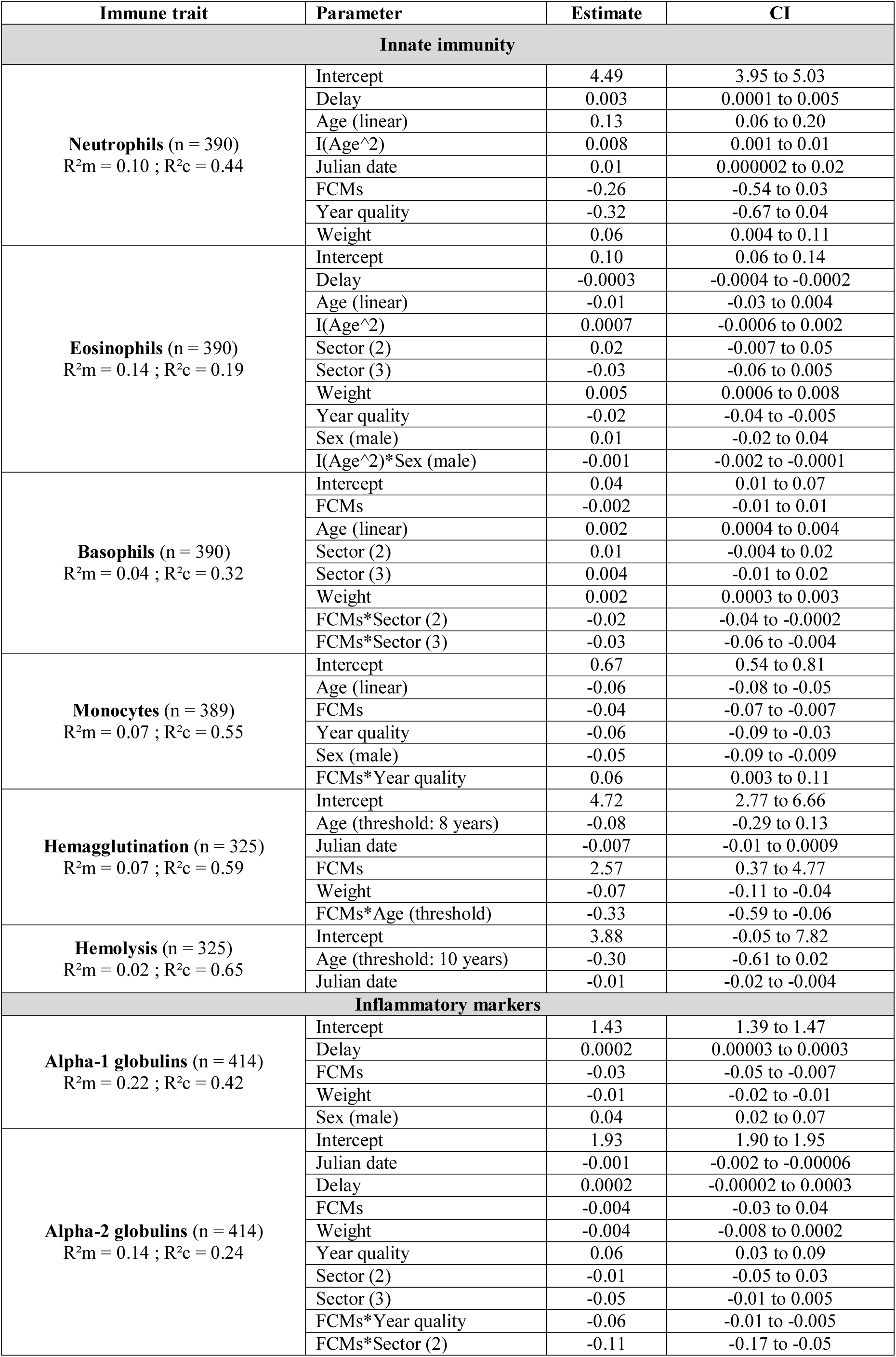
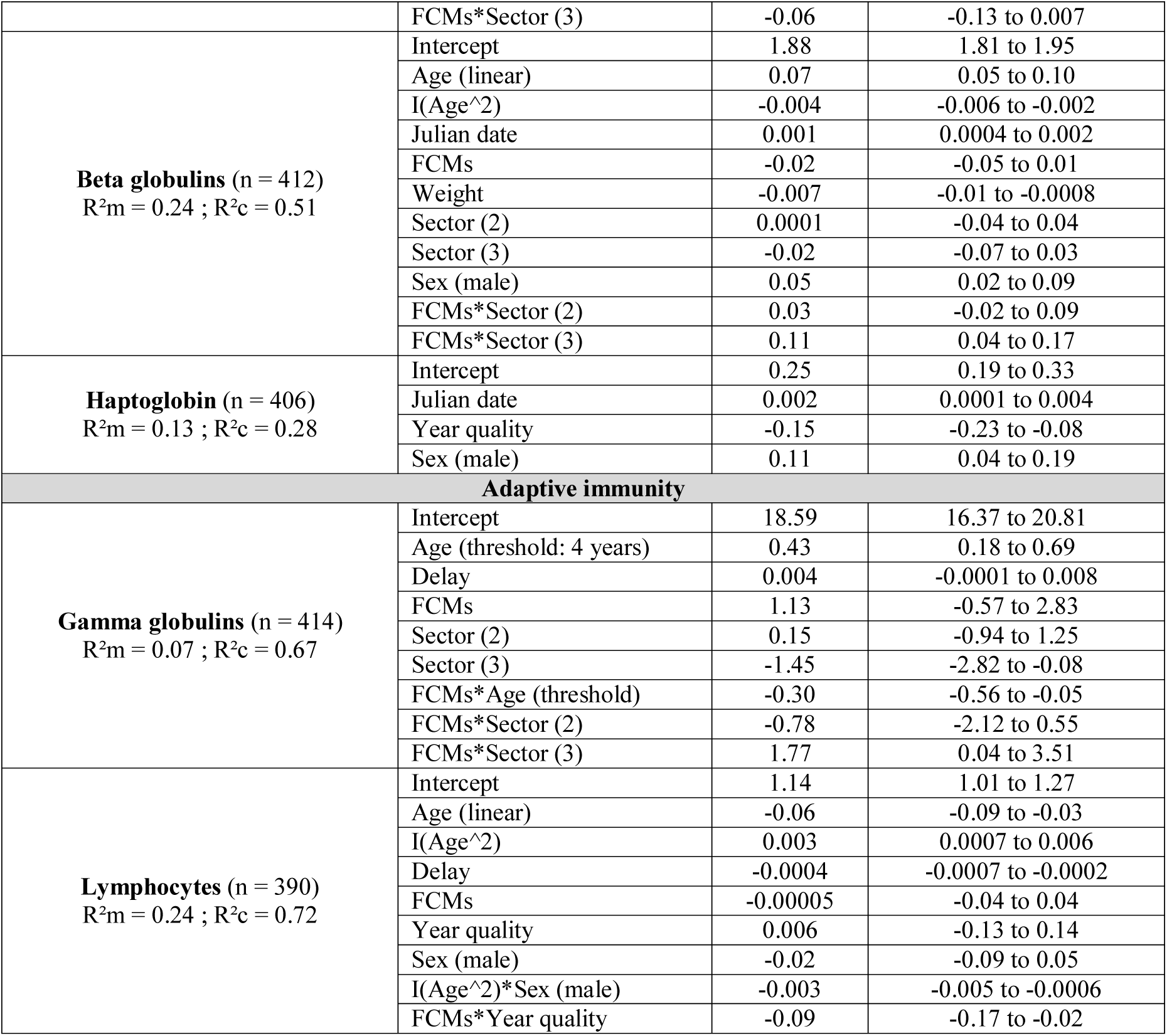
Parameters of linear mixed-effect models selected for each immune parameters of the Chizé population. R2m and R2c correspond respectively to the marginal and conditional variance explained by the model, CI corresponds to the upper and lower limits of the 95% confidence interval, and n represents the number of observations per analysis. See the material and methods section for a full definition of model sets.

**Table 3.**
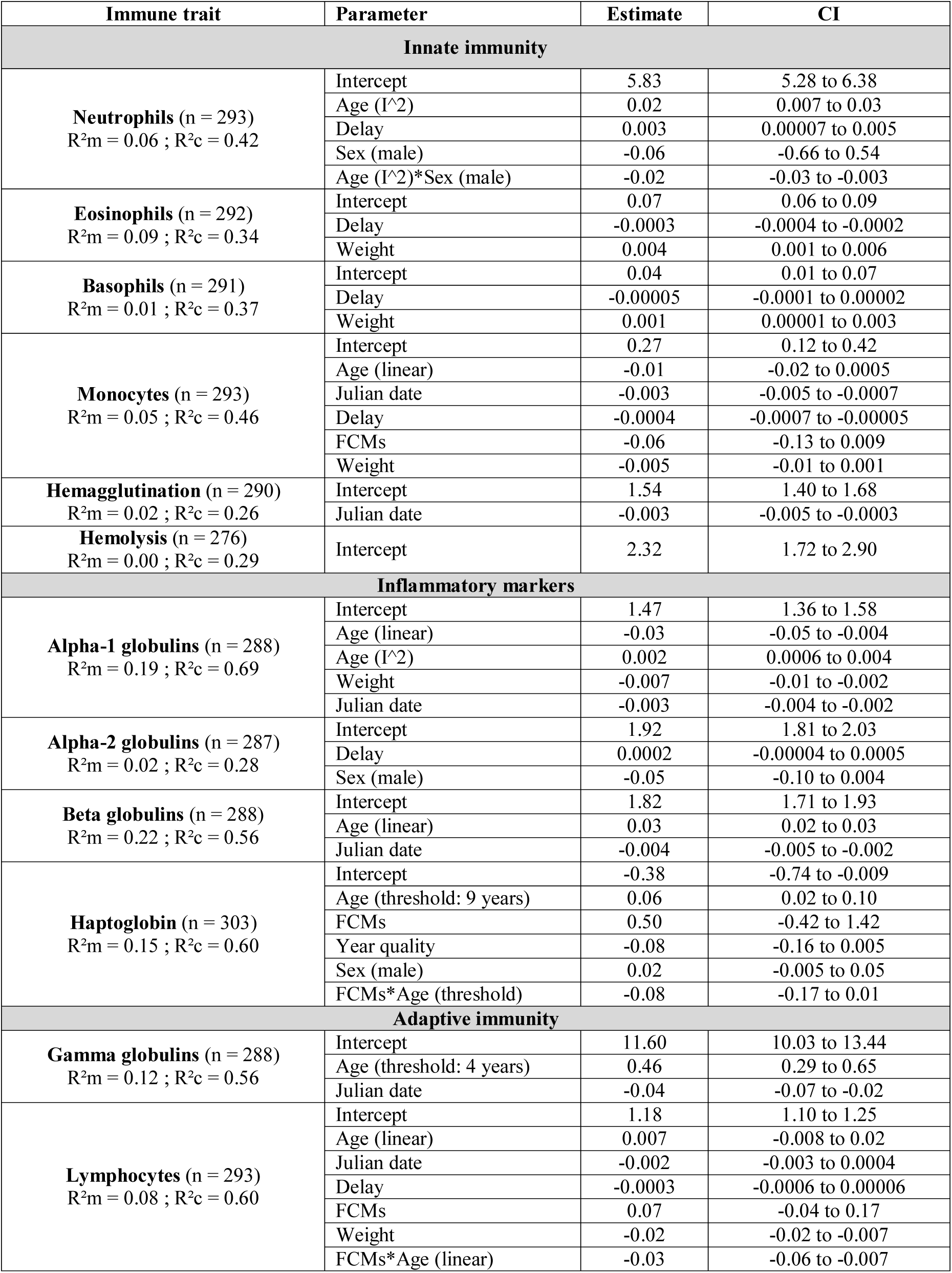
Parameters of linear mixed-effect models selected for each immune parameters of the Trois-Fontaines population. R2m and R2c correspond respectively to the marginal and conditional variance explained by the model, CI corresponds to the upper and lower limits of the 95% confidence interval, and n represents the number of observations per analysis. See the material and methods section for a full definition of model sets.

### 3.1 Innate cellular immunity

As expected from prediction 1, negative covariations between FCMs and cellular concentrations were observed for neutrophils in Aurignac (slope of -0.51; CI = [-0.94; -0.08]; Fig. 1A) with a similar trend in Chizé (−0.26; CI = [-0.54; 0.03]). A trend for a negative covariation between FCMs and monocytes was also observed in Chizé (F = 2.92; p = 0.09; slope of -0.04; CI = [-0.07 - -0.007] for average quality food resources) and in Trois-Fontaines (slope of -0.06; CI = [-0.13; 0.009]). However, in Chizé only, the negative covariation was buffered in years offering better quality food resources (Table 2; Fig. 2A), as expected from prediction 2. In Chizé, whereas basophil concentration was negatively related to FCMs levels in general (F = 6.71; p = 0.01), this particularly applied to areas of intermediate (sector 2) and poor quality (sector 3), while the covariation was close to 0 in the area of best quality (Table 2; Fig. 3A; slope of -0.002; CI = [-0.01 – 0.01]). We found no significant association between eosinophil concentrations and FCMs in any of the populations, and no interaction between FCMs and age (prediction 3) on any of the innate immune parameters measured (Table 1, 2, 3).

**Fig 1.**
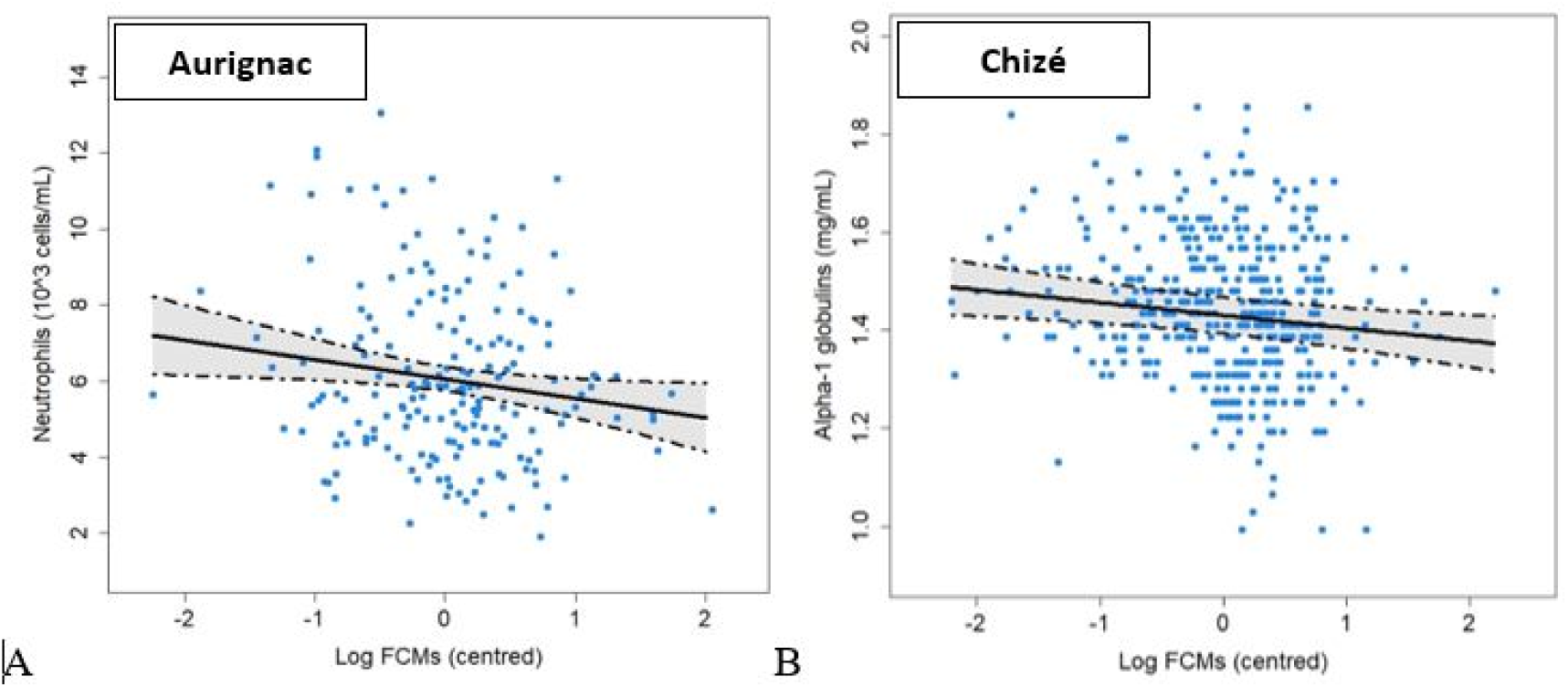
Relationship between faecal cortisol metabolites (FCMs) and immune parameters in roe deer populations (prediction (i)): A) neutrophils in Aurignac; B) alpha-1 globulins in Chizé. Points represent observed values. Solid black lines represent model predictions, dashed lines and shaded area represent 95% confidence intervals. FCMs values are centred around the mean (mean value of the variable that is subtracted from every value).

**Fig 2.**
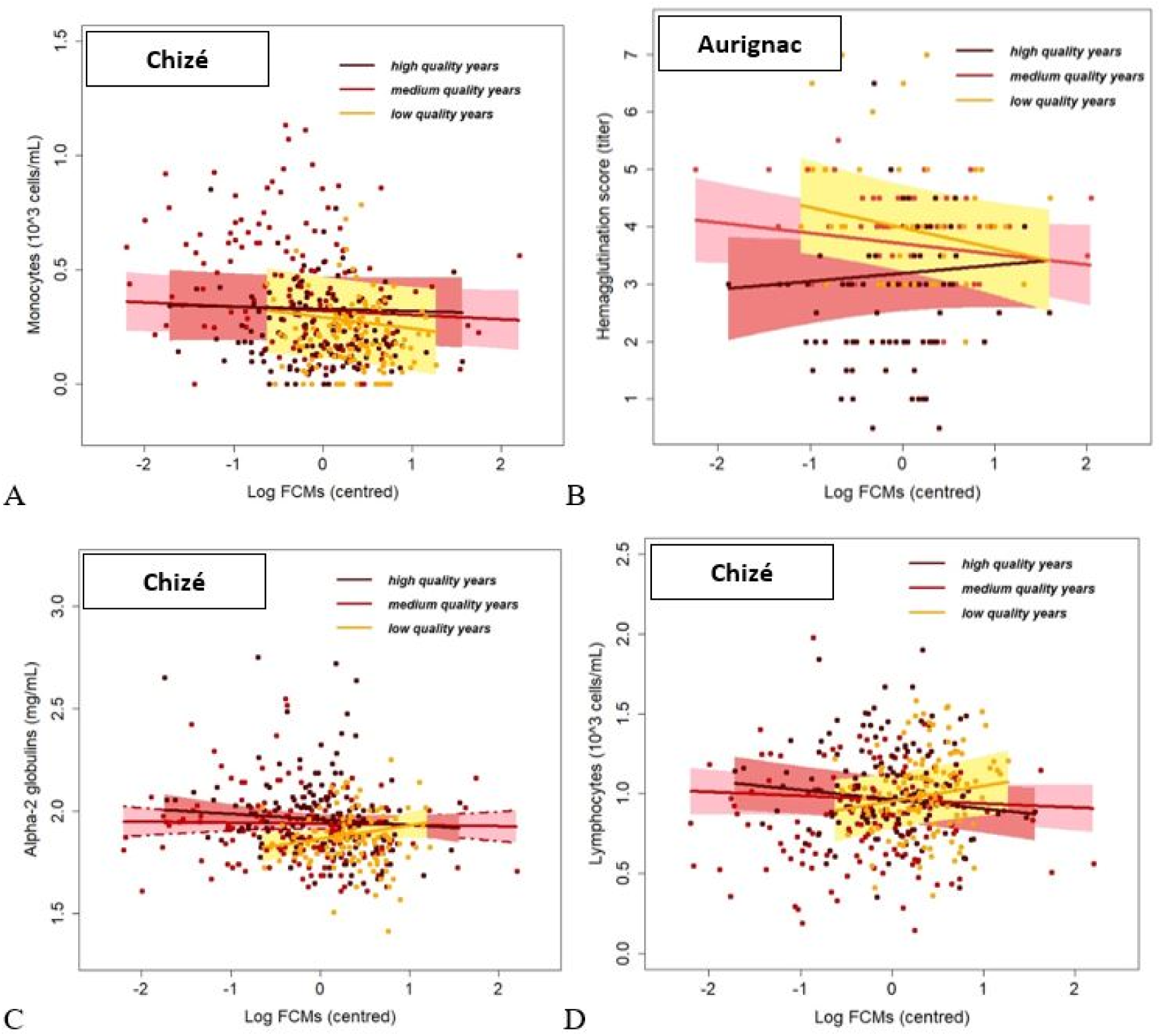
Relationship between faecal cortisol metabolites (FCMs) and immune parameters in roe deer populations at varying year quality (prediction (ii)). Year quality was indexed using the population average body mass of juveniles (in kg) captured during the following winter: A) monocytes in Chizé; B) hemaglutination score in Aurignac; C) alpha-2 globulins in Chizé; D) lymphocytes in Chizé. Points represent observed values. Solid lines represent model predictions and shaded areas represent the 95% confidence interval. FCMs values are centred around the mean (mean value of the variable that is subtracted from every value).

**Fig 3.**
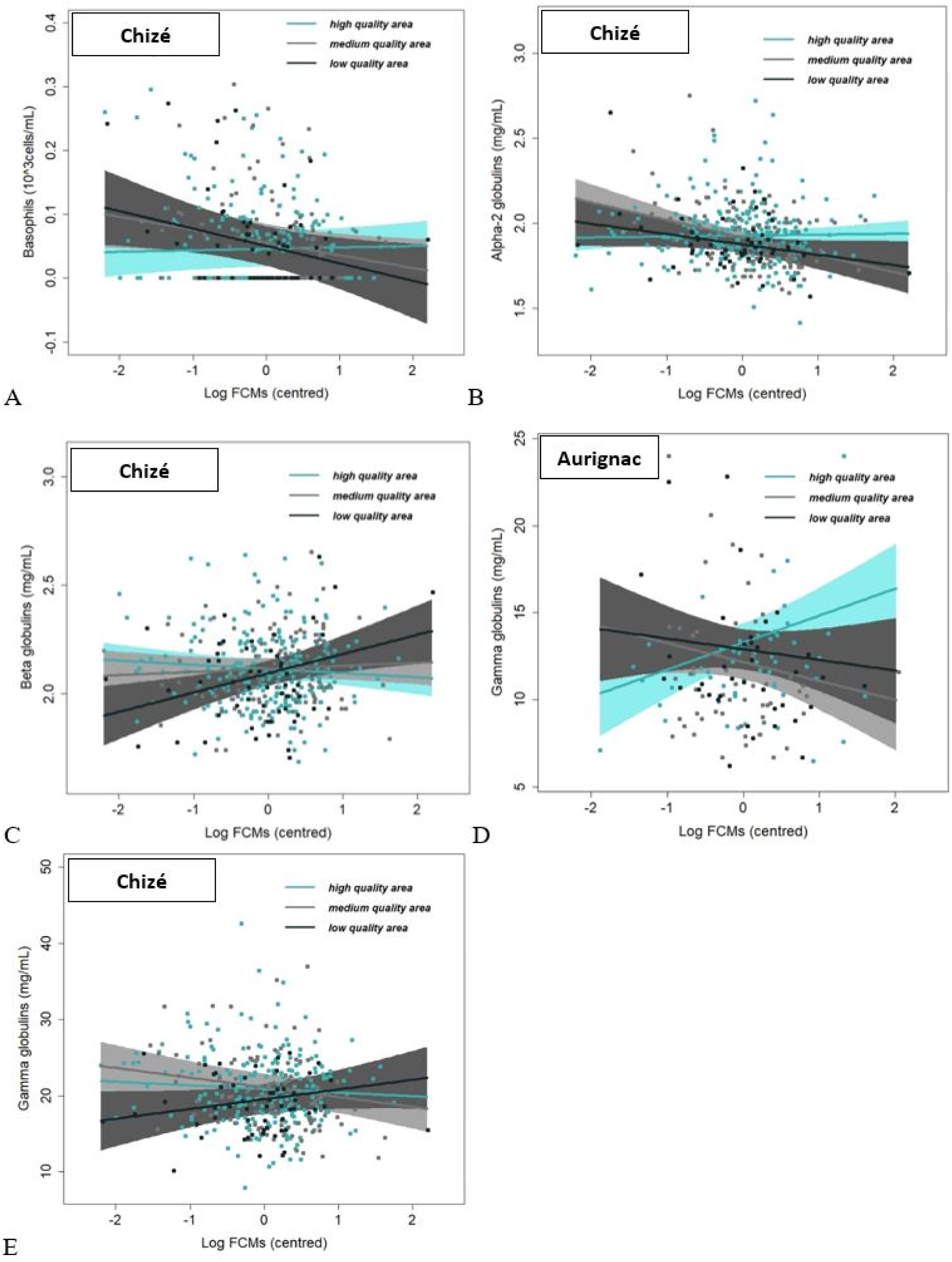
Relationship between faecal cortisol metabolites (FCMs) and immune parameters in roe deer populations, at varying area quality (prediction (ii)). Areas quality differed among the three sectors described in the material and methods section: A) basophil concentrations in Chizé; B) alpha-2 globulins in Chizé; C) beta-globulins in Chizé; D) gamma-globulins in Aurignac; E) gamma-globulins in Chizé. Points represent observed values. Solid lines represent model predictions and shaded areas represent the 95% confidence interval. FCMs values are centred around the mean (mean value of the variable that is subtracted from every value).

### 3.2 Innate humoral immunity

In Aurignac, the overall covariation between FCMs and hemagglutination titers was not significant (F = 1.69; p = 0.20; slope of -0.15; CI = [-0.37 – 0.08] for average quality resources) but a negative covariation appeared when annual food quality resources decreased, as expected from prediction 2 (Table 1; Fig 2B). In Chizé, while hemagglutination titers and FCMs were overall positively related (F = 5.80; p = 0.02; slope of 2.57; CI = [0.37 – 4.77] for animals aged less than 8 years old), the covariation became negative as age of roe deer increased (−0,33 per year; CI = [-0.59; -0.06]; Table 2; Fig. 4A). In addition, in Aurignac, FCMs were retained in the model selection to explain hemolysis titers, with a negative trend (−0.19; CI = [-0.43; 0.05]; Table 1).

**Fig 4.**
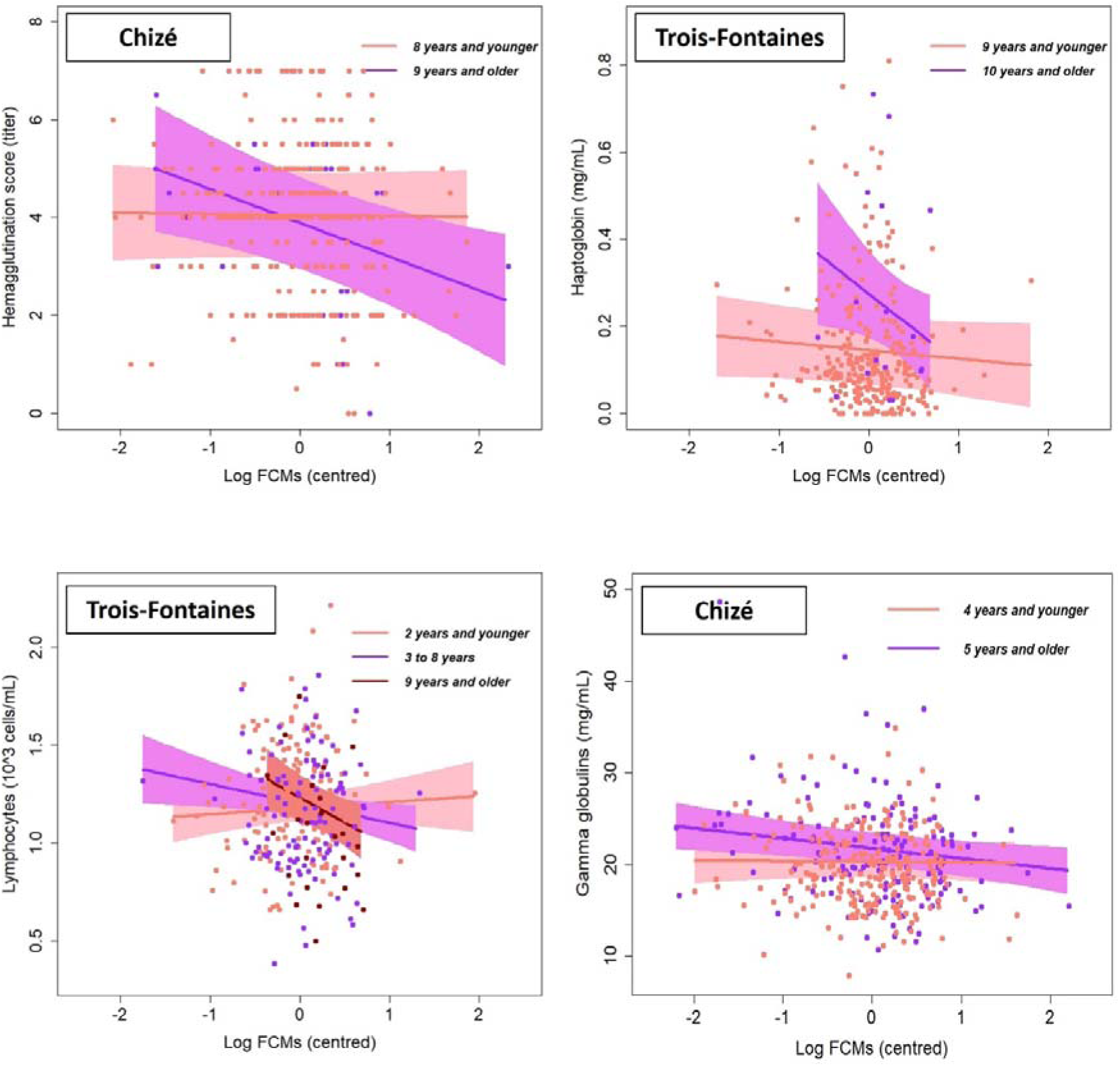
Relationship between faecal cortisol metabolites (FCMs) and immune parameters at varying ages in roe deer populations (prediction (iii)). Age was considered either as linear or with a threshold function (determined from a previous study, see main text for details), in which case the graphical representation compares before/after threshold age: A) hemagglutination in Chizé (threshold: 8 years old), B) haptoglobin in Trois-Fontaines (threshold: 9 years old); C) lymphocytes in Trois Fontaines (linear, but for the graphical display, three age classes were constituted on the basis of sample size: up to 2 years, 3-8 years, 9 years and older); D) gamma-globulins in Chizé (threshold: 4 years old). Points represent observed values. Solid lines represent model predictions and shaded areas represent the 95% confidence interval. FCMs values are centred around the mean (mean value of the variable that is subtracted from every value).

### 3.3. Inflammatory markers

In Aurignac, while alpha-1 globulin concentration was not related to FCM levels overall (F = 1.45; p = 0.23), a trend for a negative covariation appeared in areas of intermediate (sector 2) and especially poor quality (sector 3; Table 1), as expected from prediction 2. In addition, there was also a trend for a negative covariation between alpha-1 globulins and FCMs in males compared to females (slope of -0.05; CI = [-0.11; 0.01]), and in heavier individuals (slope of -0.006; CI = [-0.01; 0.002]), while beta-globulins seemed slightly positively related to FCMs (slope of 0.03; CI = [- 0.009; 0.08]). In Chizé, alpha-1 globulins were negatively related to FCMs (slope of -0.03; CI = [- 0.05; -0.007]; Table 2; Fig. 1B), as expected from prediction 1. Still in Chizé, the covariation between FCMs and alpha-2 globulins was overall negative (F = 11.9; p < 0.01), as expected from prediction 1. However, this negative covariation only held in the areas of intermediate (sector 2) and poor (sector 3) quality, while no covariation was observed in the area of best quality (Table 2; Fig. 3B). In addition, and unexpectedly, the covariation between FCMs and alpha2-globulins was more pronounced when the annual quality of food resources decreased, which is in contradiction with prediction 2 (Table 2; Fig. 2C). In Chizé again, prediction 2 was not supported by results on beta-globulin concentrations, as a positive covariation with FCMs was found in area of poor quality (sector 3), with a similar trend in the area of intermediate quality (sector 2), while no covariation was detected in the area of best quality (sectors 1, Table 2; Fig. 3C). In Trois-Fontaines, a slightly negative covariation between haptoglobin and FCM levels was observed in youngest animals but was exacerbated in old ones, in accordance with prediction 3 (slope of -0.08 per year; CI = [-0.17; - 0.01]; Table 2; Fig 4B).

### 3.4 Adaptive cellular immunity

Consistent with prediction 1, we observed no covariation between lymphocyte concentrations and FCM levels in any of the 3 populations. However, in Chizé, and contrarily to prediction 2, a positive covariation between FCMs and lymphocytes appeared during the worse quality years (Table 2; Fig. 2D). In Trois-Fontaines, a negative covariation between FCMs and lymphocytes appeared only in older roe deer (Table 3; Fig. 4C), in accordance with prediction 3.

### 3.5 Adaptive humoral immunity

Gamma-globulin concentrations were overall positively related to FCM levels in Aurignac (F = 2.62; p < 0.01), but the covariation differed among areas of different qualities, consistent with prediction 2. Precisely, the positive covariation was only present in the area of best quality (sector 1), whereas negative covariations appeared in areas of intermediate (sector 2) and poor (sector 3) quality (Table 1; Fig. 3D). Furthermore, in the Aurignac population, the covariation between gamma-globulins and FCMs differed between females and males, with females showing a positive link, while males had a negative covariation between FCMs and gamma-globulins (slope of -2.10; CI = [-3.62; -0.57]). In Chizé, a similar overall slight positive covariation was observed between gamma-globulins and FCMs (F = 3.18; p < 0.01), but the covariation became negative as age increased (slope of -0.30 per year; CI = [-0,56; -0.05]; Table 2; Fig. 4D). As in Aurignac, the positive covariation between gamma-globulin concentrations and FCMs also varied depending on the sector. However, contrary to Aurignac and to prediction 2, in Chizé, the covariation was positive only in the area of poor quality (Table 2; Fig. 3E). No significant correlation with FCMs was observed in the Trois-Fontaines population.

## 4. Discussion

Overall, our results highlight clear associations between baseline glucocorticoid levels and immune parameters and show that the strength and direction of these associations differ between the three studied populations. Of the 12 measured immune parameters, only three (monocytes, lymphocytes and haptoglobin) showed significant co-variations with FCMs in Trois-Fontaines, and six (neutrophils, hemagglutination, hemolysis, alpha-1, beta and gamma-globulins) in Aurignac. The co-variation between FCMs and immunological parameters was particularly marked in Chizé (nine parameters), a site where the population faces harsh environmental conditions (Gaillard et al., 1993). Taken together, our results constitute one of the rare pieces of evidence that adverse effects on immune functions due to long-term elevation of glucocorticoid levels (Dhabhar and McEwen, 1997; Sapolsky et al., 2000; Dhabhar, 2009; Stier et al., 2009) also occur in the wild.

Furthermore, and as expected, we found that covariations between FCMs and immune parameters differed between immunity components and that these covariations were affected by individual and environmental factors. Our results support the hypothesis that immune system components are differentially affected by glucocorticoid levels (Bourgeon and Raclot, 2006). As levels of FCMs increased, we observed an overall decline in cellular (neutrophils, monocytes, basophils) and humoral (hemolysis) innate immune functions, in accordance with prediction 1 and previous reports. For instance, in the Belding’s ground squirrel (*Urocitellus beldingi*), experimental chronic elevation of cortisol levels reduced serum bactericidal competence, a component of the constitutive innate immune response, compared to a control group (Brooks and Mateo, 2013). The most plausible explanation for this negative covariation between FCMs and innate immune parameters relies on the high energy cost of this immune system component. Indeed, the maintenance of immune defences requires energy and nutrients (Lochmiller and Deerenberg, 2000), and the cost of activation is higher for innate than for adaptive immunity (McDade, 2016). Consequently, long-term elevation of glucocorticoid levels may selectively redirect energy away from the innate immunity towards other energy demanding functions (Lee, 2006; Martin, 2009). This process could be seen as a physiological adjustment for energy savings to maximize investment in reproduction and long-term survival through mechanisms other than innate immunity.

Our results show that, of the four inflammatory markers measured, three (alpha-1, alpha-2 and haptoglobin) were negatively associated with FCMs in line with our predictions. Indeed, negative covariations between FCMs and inflammatory markers are consistent with the energy trade-off hypothesis, and likely due to their particularly high activation costs (McDade, 2016). However, we also found weak positive covariations between FGMs and beta-globulins. Why the relationship was positive for one of the inflammatory markers while it was negative for the others remains unanswered. First, beta-globulins, like alpha-1 and alpha-2 globulins, are a complex group of proteins, that may each have distinct relationship with stress level. Previous studies also pointed out that the relationship between glucocorticoids and immune parameters of the same arm may be different, such that some antigen responses in chickens have been shown to be affected by stressors, while others were not (El-Lethey et al., 2003). The proposed explanation for these differences was an insensitivity to glucocorticoids for certain immune parameters, which could be a protective mechanism. The discrepancy between markers of the same arm of immunity highlights an important aspect of our work: the complexity of the relationship between immunity and glucocorticoids calls for great caution in the choice of the immune markers and in their interpretations.

Our results also partially supported our prediction of a moderate or absence of covariation between FCMs and cellular (lymphocytes) or humoral (gamma-globulins) adaptive immune parameters, which we observed in the three studied populations. Since the activation costs of adaptive immunity are lower than those of innate immunity (McDade, 2016), adaptive immune functions are expected to be less prone to energy trade-off. An alternative non-exclusive explanation to this lack of link, or positive covariation (in the case of gamma-globulins in Aurignac in the high-quality sector), between FCMs and adaptive immune parameters may be that individuals repeatedly or chronically exposed to stressors may actually allocate more in adaptive immunity, especially if major stressors are pathogens, in order to maximize long-term survival. This might be particularly true in long-lived species such as roe deer, which repeatedly face the same pathogens in their environment, and are expected to exhibit stronger allocation in adaptive, and particularly antibody-mediated immunity, compared to innate immunity (Lee, 2006).

Besides the difficulty of making general assumptions about the immunity-glucocorticoid relationship due to the above-mentioned differences between components, we found that other factors may modulate the observed relationships. In particular, the age of individuals as well as the spatial and annual heterogeneity of food resources influenced most of the covariations between FCMs and innate, inflammatory, and adaptive markers of immunity in the three studied populations. As predicted in 2, we found that during years providing less and/or lower-quality food resources a negative covariation between innate immunity (monocytes and hemagglutination) and baseline glucocorticoid levels appeared, contrary to better years. The same observation was made for the spatial heterogeneity in food resources. In the Chizé and Aurignac populations, roe deer from areas with low quality food resources showed negative covariations between some immune markers (basophils, alpha-1, alpha-2 and gamma globulins) and FCM levels, while no or positive covariations were observed in areas of high quality. High levels of food resources have been shown to be associated with reduced glucocorticoid levels (Fokidis et al., 2012; Carbillet et al., 2020) but may also provide sufficient energy and nutrients to sustain the cost of innate immune response, as previously suggested in by Strandin and his colleagues (2018). In their review, these authors showed that food provisioning in field studies tended to increase both innate and adaptive immunity, whereas food restriction frequently impaired immunity. However, in contrast to those results, our data also suggested that negative covariations may, in some cases, appear between immunity and baseline glucocorticoid levels in areas and years of high food resources. Alpha-2 globulins and lymphocytes were positively associated with FCM levels when annual food resources were scarce, and there were positive covariations with FCMs for beta-globulins and gamma globulins in poor food resources areas, while no association was observed in areas providing better food resources. At first glance, these results might appear counter-intuitive. However, these observations were all made in the population of Chizé, which has the lowest availability in food resources compared to the two other populations, even in the best sector (Gaillard et al., 1993). These results thus seem to match with an alternative theoretical framework proposed by Davis and Maney (2018), who suggested that in poor quality habitats, glucocorticoid secretion should be downregulated to prevent impairment of immune functions. Therefore, in Chizé, during the worst years, or in the worst sector, energy allocation to immune function could be prioritised and independent of glucocorticoid levels, which should be minimised with low among-individual variations. Consistent with this hypothesis, variance in FCMs appeared to be lower during years of low food availability than during years of high food availability in Chizé (Fig.2). However, this hypothesis is not supported for all immune parameters studied, suggesting that not all immune parameters would be independent of glucocorticoid levels in poor quality habitats within the framework proposed by Davis and Maney (2018). In addition, pathogen load, and thus immune challenge, may covary with food resources availability across time. In particular, variations in population abundance may determine both resource availability and pathogen exposure: high population density leading to both limited resources and high exposure to directly or indirectly transmitted pathogens. In this case, high pathogen pressure could act as a stressor on roe deer and lead to an elevation of FCMs, while stimulating immune defences to cope with the threat.

Finally, age influenced the relationship between immunity and baseline glucocorticoid levels. In accordance with our predictions 3, negative relationships between FCMs and immune parameters (hemagglutination, haptoglobin, gamma globulins and lymphocytes) only appeared, or were steeper, in older individuals. In roe deer, the ingestion capacity is known to be less efficient in old individuals (Gaillard et al., 1993), leading to a diminution in available energy and nutrients, and individuals suffer from a loss of condition, as documented through a senescence in body mass (Douhard et al. 2017) and various biological markers of ageing (Cheynel et al. 2017, Wilbourn et al. 2017; Lemaître et al. 2022). Consequently, older individuals exhibiting high levels of baseline glucocorticoids might have poorer abilities to increase their allocation toward the immune functions. Moreover, in line with previous evidence that stress hormones might accelerate the aging process in roe deer (Lemaître et al. 2021), our results suggest that they might also accelerate the previously documented immunosenescence (Cheynel et al. 2017). Long-term or repeated exposure to stressors may therefore constitute a major selective pressure, especially on old individuals that may not be able to maintain an overall efficient immune response and would therefore be exposed to higher risks of diseases, contributing to the reduction in survival and reproductive success.

To conclude, our results show that the immunity of wild ungulates is strongly shaped by baseline glucocorticoid levels, but that this influence differ between innate, adaptive and inflammatory markers of immunity. While glucocorticoids are overall negatively correlated with innate and inflammatory immunity, they appear to be less or even positively linked to adaptive immunity. In addition, spatial and temporal availability in food resources appear to shape the relationship between glucocorticoids and immunity with non-linear patterns, while negative covariations between FCMs and immunity is strongest in old individuals. Our work highlights the need to consider a multi-marker approach for future studies investigating the effect of stress hormones on immune functions in wild animals. Such an approach, combined with consideration of the environmental context and individual phenotype, could help improve our understanding of the mechanisms underlying among-individual differences in immunity and susceptibility to diseases in the wild.

## Supporting information

Model selection Chize

Model selection Trois Fontaines

Model selection Aurignac

Correlation matrix

Predictions

## Acknowledgments

We thank all the LBBE, CEFS, ONCFS/OFB staff and all the field volunteers for the organisation and their assistance during the roe deer captures. We thank the local hunting associations and the Fédération Départementale des Chasseurs de la Haute Garonne. We also thank Edith Klobetz-Rassam for EIA analysis.

## Authors’ contributions

JC, EGF and HV conceived and designed the study. JC, EGF, HV, BR, MP, JD, SP, FD, JM, JFL performed fieldwork. EGF, AG and CR performed immunological analyses. RP and JC ran FCM assays. JC and MH performed the statistical analysis, wrote the first draft of the paper, and then received input from other co-authors. All authors approved the final version of the manuscript and agree to be held accountable for the content therein.

## Data accessibility

Data used in this study are available at https://github.com/JeffreyCarbillet/CarbilletHollainEtal2022

## References

Abbas, F., Morellet, N., Hewison, A.M., Merlet, J., Cargnelutti, B., Lourtet, B., Angibault, J.M., Daufresne, T., Aulagnier, S., Verheyden, H., 2011 Landscape fragmentation generates spatial variation of diet composition and quality in a generalist herbivore. Oecologia, 167: 401–411. 10.1007/s00442-011-1994-0 (doi.org)

Arnold, T. W., (2010) Uninformative parameters and model selection using Akaike’s Information Criterion. J. Wildl. Manage. 74: 1175–1178. https://doi.org/10.1111/j.1937-2817.2010.tb01236.x

Barton, K., 2016. MuMIn: multi-model inference. R package version 1.15.6 Available from https://cran.r-project.org/web/packages/MuMIn/index.html.

Bates, D., Mächler, M., Bolker, B., Walker, S., 2014. Fitting linear mixed-effects models using lme4. Preprint. Available from: arXiv:1406.5823.

Bonnot, N., Morellet, N., Verheyden, H., Cargnelutti, B., Lourtet, B., Klein, F., Hewison, A.J.M., 2013. Habitat use under predation risk : hunting, roads and human dwellings influence the spatial behaviour of roe circles. Eur. J. Wildl. Res. 59: 185–193. https://doi.org/10.1007/s10344-012-0665-8

Boonstra, R., 2005. Equipped for life: the adaptive role of the stress axis in male mammals. J. Mammal. 86: 236–247. https://doi.org/10.1644/BHE-001.1

Bourgeon, S., Raclot, T., 2006. Corticosterone selectively decreases humoral immunity in female eiders during incubation. J. Exp. Biol. 209: 4957–4965. https://doi.org/10.1242/jeb.02610

Breuner, C.W., Patterson, S.H., Hahn, T.P., 2008. In search of relationships between the acute adrenocortical response and fitness. Gen. Comp. Endocrinol. 157: 288–295. https://doi.org/10.1016/j.ygcen.2008.05.017

Brooks, K.C., Mateo, J.M., 2013. Chronically raised glucocorticoids reduce innate immune function in Belding’s ground squirrels (*Urocitellus beldingi*) after an immune challenge. Gen. Comp. Endocrinol. 193: 149–157 https://doi.org/10.1016/j.ygcen.2013.07.019

Burnham, K.P., Anderson, D.R., (2003) Model selection and multimodel inference: a practical information-theoretic approach. Springer Science & Business Media. https://doi.org/10.1007/b97636

Carbillet, J., Rey, B., Palme, R., Morellet, N., Bonnot, N., Chaval, Y., Cargnelutti, B., Hewison, A.J.M., Gilot-Fromont, E. and Verheyden, H., 2020. Under cover of the night: context-dependency of anthropogenic disturbance on stress levels of wild roe deer *Capreolus capreolus*. Conserv. Physiol. 8: coaa086. https://doi.org/10.1093/conphys/coaa086

Carbillet, J., Rey, b., Lavabre, T., Geffre, A., Chaval, Y., Merlet, J., Débias, F., Régis, C., Pardonnet, S., Gaillard, J.M., Hewison, A.J.M., Lemaître, J.F., Pellerin, M., Rannou, B., Verheyden, H., Gilot-Fromont, E., 2019. Using the Neutrophil to Lymphocyte ratio to highlight individual coping styles across variable environments in the wild. Behav. Ecol. Sociobiol. 73: 144 https://doi.org/10.1007/s00265-019-2755-z

Cheynel, L., Lemaître, J.F., Gaillard, J.M., Rey, B., Bourgoin, G., Ferté, H., Jégo, M., Débias, F., Pellerin, M., Jacob, L., Gilot-Fromont, E., 2017. Immunosenescence patterns differ between populations but not between sexes in a long-lived mammal. Sci. Rep. 7: 1–11. https://doi.org/10.1038/s41598-017-13686-5

Davis, A.K., Maney, D.L., 2018. The use of glucocorticoid hormones or leucocyte profiles to measure stress in vertebrates: What’s the difference? Methods Ecol. Evol. 9: 1556–1568. https://doi.org/10.1111/2041-210X.13020

Dhabhar, F.S., Malarkey, W.B, Neri, E., McEwen, B.S. 2012. Stress-induced redistribution of immune cells–from barracks to boulevards to battlefields: a tale of three hormones-curt Richter Award winner. Psychoneurendocrinology 37: 1345–1368. https://doi.org/10.1016/j.psyneuen.2012.05.008

Dhabhar, F.S., 2009. Enhancing versus suppressive effects of stress on immune function: implications for immunoprotection and immunopathology. Neuroimmunomodulation 16: 300–317. https://doi.org/10.1159/000216188

Dhabhar, F.S., McEwen, B.S., 1997. Acute stress enhances while chronic stress suppresses cell-mediated immunity in vivo: a potential role for leukocyte trafficking. Brain Behav. Immun. 11: 286–306. https://doi.org/10.1006/brbi.1997.0508

Dhabhar, F.S., Miller, A.H., McEwen, B.S., Spencer, R.L., 1995. Effects of Stress on Immune Cell Distribution. J. Immunol. 154: 5511–5527. https://doi.org/10.1016/B978-012373947-6.00217-8

Dhabhar, F.S., Miller, A.H., Stein, M., McEwen, B.S., Spencer, R.L., 1994. Diurnal and acute stress-induced changes in distribution of peripheral blood leukocyte subpopulations. Brain Behav. Immun. 8: 66–79. https://doi.org/10.1006/brbi.1994.1006

Douhard, M., Plard, F., Gaillard, J.M., Capron, G., Delorme, D., Klein, F., Duncan, P., Loe, L.E., Bonenfant, C., 2014. Fitness consequences of environmental conditions at different life stages in a long-lived vertebrate. Proc Royal Soc. B. 281: 20140276. https://doi.org/10.1098/rspb.2014.0276

Douhard, F., Gaillard, J.M., Pellerin, M., Jacob, L. and Lemaître, J.F., 2017. The cost of growing large: Costs of post□weaning growth on body mass senescence in a wild mammal. Oikos. 126: 1329–1338. https://doi.org/10.1111/oik.04421

El-Lethey, H., Huber-Eicher, B., & Jungi, T. W., 2003. Exploration of stress-induced immunosuppression in chickens reveals both stress-resistant and stress-susceptible antigen responses. Vet. Immunol. Immunopathol., 95: 91–101. https://doi.org/10.1016/S0165-2427(02)00308-2

Fokidis, H.B., Des Roziers, M.B., Sparr, R., Rogowski, C., Sweazeazea, K., Deviche, P., 2012. Unpredictable food availability induces metabolic and hormonal changes independent of food intake in a sedentary songbird. J. Exp. Biol. 215: 2920–2930. https://doi.org/10.1242/jeb.071043

Gaillard, J.M., Delorme, D, Boutin, J.M., Van Laere, G, Boisaubert, B., Pradel, R., 1993. Roe deer survival patterns – a comparative analysis of contrasting populations. J. Anim. Ecol. 62: 778–791. https://doi.org/10.2307/5396

Gilot-Fromont, E., Jégo, M., Bonenfant, C., Gilbert, P., Rannou, B., Klein, F., Gaillard, J.M., 2012. Immune Phenotype and Body Condition in Roe Deer: Individuals with High Body Condition Have Different, Not Stronger Immunity. PLoS One 7: e45576. https://doi.org/10.1371/journal.pone.0045576

Hau, M., Casagrande, S., Ouyang, J.Q., Baugh, A.T., 2016. Glucocorticoid-Mediated Phenotypes in Vertebrates: Multilevel Variation and Evolution. Adv. Study Behav. 48: 41–115. https://doi.org/10.1016/bs.asb.2016.01.002

Hewison, A.J.M., Morellet, N., Verheyden, H., Daufresne, T., Angibault, J-M., Cargnelutti, B., Merlet, J., Picot, D., Rames, J-L., Joachim, J., Lourtet, B., Serrano, E., Bideau, E., Cebe, N., 2009. Landscape fragmentation influences winter body mass of roe deer. Ecography 32: 1062–1070. https://doi.org/10.1111/j.1600-0587.2009.05888.x

Hewison, A.J.M., Angibault, J.M., Cargnelutti, B., Coulon, A., Rames, J.L., Morellet, N., Verheyden, H., Serrano, E., 2007. Using Radio-tracking and Direct Observation to Estimate Roe Deer Capreolus Capreolus Density in a Fragmented Landscape: A Pilot Study. Wildlife Biol. 13: 313–320. https://doi.org/10.2981/0909-6396(2007)13[313:URADOT]2.0.CO;2

Hewison, A.J.M., Vincent, J.P., Angibault, J.M., Delorme, J., Van Laere, G., Galliard J.M., 1999. Tests of estimation of age from tooth wear on roe deer of known age: variation within and among populations. Can. J. Zool. 77: 58–67. https://doi.org/10.1139/z98-183

Houwen, B., (2001). The differential cell count. Int. J. Lab. Hematol. 7: 89–100.

Hurlbert, S.H., 1984. Pseudoreplication and the design of ecological field experiments. Ecol. Monogr. 54: 187–211. https://doi.org/10.2307/1942661

Jain, S., Gautam, V., and Naseem, S., 2011. Acute-phase proteins: As diagnostic tool. J. Pharm. Bioallied Sci. 3: 118.

Josserand, R., Haussy, C., Agostini, S., Decencière, B., Le Galliard, J. F., & Meylan, S., (2020). Chronic elevation of glucorticoids late in life generates long lasting changes in physiological state without a life history switch. Gen. Comp. Endocrinol. 285: 113288. https://doi.org/10.1016/j.ygcen.2019.113288

Kirkwood, T.B.L., 1977. Evolution of ageing. Nature 270: 301–304. https://doi.org/10.1038/270301a0

Klasing, K.C., Leshchinsky, T.V., 1999. Functions, costs, and benefits of the immune system during development and growth. Ostrich, 69: 2817–32.

Klasing, K.C., 2004. The cost of immunity. Acta Zoologica Sinica, 50: 961–969.

Lee, K.A., 2006. Linking immune defenses and life history at the levels of the individual and the species. Integr. Comp. Biol. 46: 1000–1015. https://doi.org/10.1093/icb/icl049

Lemai□tre, J.F., Carbillet, J., Rey, B., Palme, R., Froy, H., Wilbourn, R.V., Underwood, S.L., Cheynel, L., Gaillard, J.M., Hewison, A.M. and Verheyden, H., 2021. Short-term telomere dynamics is associated with glucocorticoid levels in wild populations of roe deer. Comp. Biochem. Physiol. Part A Mol. Integr. Physiol. 252: 110836. https://doi.org/10.1016/j.cbpa.2020.110836

Lemaître, J.F., Rey, B., Gaillard, J.M., Régis, C., Gilot, E., Debias, F., Duhayer, J., Pardonnet, S., Pellerin, M., Haghani, A., Zoller, J.A., Li, C.Z. and Horvath, S., 2022. DNA methylation as a tool to explore ageing in wild roe deer populations. Molecular ecology resources. Mol. Ecol. https://doi.org/10.1111/1755-0998.13533

Lochmiller, R.L., Deerenberg, C., 2000. Trade-offs in evolutionary immunology: just what is the cost of immunity? Oikos. 88 : 87–98. https://doi.org/10.1034/j.1600-0706.2000.880110.x

Martin, J., Vourc’h, G., Bonnot, N., Cargnelutti, B., Chaval, Y., Lourtet, B., Goulard, M., Hoch, T., Plantard, O., Hewison, A.J.M., Morellet, N., 2018. Temporal shifts in landscape connectivity for an ecosystem engineer, the roe deer, across a multiple-use landscape. Landsc. Ecol. 33: 937–954. https://doi.org/10.1007/s10980-018-0641-0

Martin, L.B., 2009. Stress and immunity in wild vertebrates: Timing is everything. Gen. Comp. Endocrinol. 163: 70–79. https://doi.org/10.1016/j.ygcen.2009.03.008

Matson, K. D., Robert, E., Ricklefs, R. E., Klasing, K.C., 2005. A hemolysis–hemagglutination assay for characterizing constitutive innate humoral immunity in wild and domestic birds. Dev. Comp. Immunol. 29: 275–286. https://doi.org/10.1016/j.dci.2004.07.006

McDade, T.W., Georgiev, A.V., Kuzawa, C.W., 2016. Trade-offs between acquired and innate immune defenses in humans. Evol. Med. Public Health 1: 1–16. https://doi.org/10.1093/emph/eov033

Metcalf, C.J.E., Roth, O. and Graham, A.L., 2020. Why leveraging sex differences in immune trade□offs may illuminate the evolution of senescence. Funct. Ecol., 34: 129–140. https://doi.org/10.1111/1365-2435.13458

Moazzam, S., Hussain, M.M., Saleem, S., 2012. Effect of ascorbic acid and alpha tocopherol on immune status of male Sprague Dawley rats exposed to chronic restraint stress. JAMC 24: 31–35.

Morellet, N., Verheyden, H., Angibault, J.M., Cargnelutti, B., Lourtet, B., Hewison, A.J.M. 2009. The effect of capture on ranging behaviour and activity of the European roe cerf *Capreolus capreolus*. Wildlife Biol., 15: 278–287. https://doi.org/10.2981/08-084

Möstl, E., Palme, R., 2002. Hormones as indicators of stress. Domest. Anim. Endocrinol. 23: 67–74. https://doi.org/10.1016/S0739-7240(02)00146-7

Möstl, E., Maggs, J.L., Schrötter, G., Besenfelder, U., Palme, R. 2002. Measurement of cortisol metabolites in faeces of ruminants. Vet. Res. Commun. 26: 127–139. https://doi.org/10.1023/A:1014095618125

Nussey D.H., Watt K., Pilkington J.G., Zamoyska R., McNeilly T.N., 2012. Age-related variation in immunity in a wild mammal population. Aging Cell 11: 178–180. https://doi.org/10.1111/j.1474-9726.2011.00771.x

Palme, R., 2019. Non-invasive measurement of glucocorticoids: advances and problems. Physiol. Beh. 199: 229–243. https://doi.org/10.1016/j.physbeh.2018.11.021

Palme, R., Touma, C., Arias, N., Dominchin, M.F., Lepschy, M., 2013. Steroid extraction: Get the best out of faecal samples. Wiener Tierärztl. Monatsschr. 100: 238–246.

Palme, R., Rettenbacher, S., Touma, C., El-Bahr, S.M., Möstl, E., 2005. Stress hormones in mammals and birds: comparative aspects regarding metabolism, excretion, and non-invasive measurement in fecal samples. Ann. N. Y. Acad. Sci. 1040: 162–171. https://doi.org/10.1196/annals.1327.021

Pettorelli, N., Gaillard, J.M., Mysterud, A., Duncan, P., Stenseth, N.C., Delorme, D., Van Laere, G., Toïgo, C., Klein, F., Ranta, E., 2006. Using a proxy of plant productivity (NDVI) to find key periods for animal performance: the case of roe deer. Oikos 112: 565–572. https://doi.org/10.1111/j.0030-1299.2006.14447.x

Pettorelli, N., Dray, S., Gaillard, J.M., Chessel, D., Duncan, P., Illius, A., Guillon, N., Klein, F., Van Laere, G., 2003. Spatial variation in springtime food resources influences the winter body mass of roe deer fawn. Oecologia, 137: 363–369. https://doi.org/10.1007/s00442-003-1364-7

Pettorelli, N., Gaillard, J.M., Duncan, P., Ouellet, J.P., Van Laere, G., 2001. Population density and small-scale variation in habitat quality affect phenotypic quality in roe circles. Oecologia, vol. 128: 400–405. https://doi.org/10.1007/s004420100682

Richards, S.A., (2008) Dealing with overdispersed count data in applied ecology. J. Appl. Ecol. 45: 218–227. https://doi.org/10.1111/j.1365-2664.2007.01377.x

Rolff J., 2002. Bateman’s principle and immunity. Proc. Royal Soc. B. 269: 867–872. https://doi.org/10.1098/rspb.2002.1959

Romero, L.M., 2004. Physiological stress in ecology: lessons from biomedical research. TREE 19: 249–255. https://doi.org/10.1016/j.tree.2004.03.008

Sapolsky, R. M., Romero, L.M., Munck, A.U., 2000. How Do Glucocorticoids Influence Stress Responses? Integrating Permissive, Suppressive, Stimulatory, and Preparative Actions. Endocr. Rev., 21: 55–89. https://doi.org/10.1210/edrv.21.1.0389

Sapolsky, R. M., Krey, L. C., McEwen, B. S., 1983. The adrenocorticol stress-response in the aged male rat: impairment of recovery from stress. Exp. Gerontol. 18: 55–64. https://doi.org/10.1016/0531-5565(83)90051-7

Sheriff, M.J., Dantzer, B., Delehanty, B., Palme, R., Boonstra, R., 2011. Measuring stress in wildlife: techniques for quantifying glucocorticoids. Oecologia 166: 869–887. https://doi.org/10.1007/s00442-011-1943-y

Sorci, G., Boulinier, T., Gauthier-Clerc, M., Faivre, B., 2008. The evolutionary ecology of the immune response. In: Thomas, F., Guégan, J.F., Renaud, F. Ecology and Evolution of Parasitism, Oxford University Press, pp. 5–17.

Stanley J., 2002. Essentials of immunology and serology. Albany, NY: Thomson Delmar Learning.

Stier, K.S., Almasi, B., Gasparini, J., Piault, R., Roulin, A., Jenni, L., 2009. Effects of corticosterone on innate and humoral immune functions and oxidative stress in barn owl nestlings. J. Exp. Biol. 212: 2085–2091. https://doi.org/10.1242/jeb.024406

Stockham S.L. & Scott M.A., (2008) Fundamentals of veterinary Clinical Pathology, second ed. Blackwell Publishing, Ames, IA.

Strandin, T., Babayan, S.A., and Forbes, K.M., 2018. Reviewing the effects of food provisioning on wildlife immunity. Proc Royal Soc. B. 373 : 20170088. https://doi.org/10.1098/rstb.2017.0088

Wada, T., Ishiwata, K., Koseki, H., Ishikura, T., Ugajin, T., Ohnuma, N., Obata, K., Ishikawa, R., Yoshikawa, S., Mukai, K., Kawano, Y., Minegishi, Y., Yokozeki, H., Watanabe, N., Karasuyama, H., 2010. Selective ablation of basophils in mice reveals their nonredundant role in acquired immunity against ticks. J. Clin. Investig. 120: 2867–2875. https://doi.org/10.1172/JCI42680

Wilbourn, R.V., Froy, H., McManus, M.C., Cheynel, L., Gaillard, J.M., Gilot-Fromont, E., Regis, C., Rey, B., Pellerin, M., Lemaître, J.F. and Nussey, D.H., 2017. Age-dependent associations between telomere length and environmental conditions in roe deer. Biol. Lett., 13: 20170434. https://doi.org/10.1098/rsbl.2017.0434

Wingfield, J. C., Sapolsky, R. M., 2003. Reproduction and resistance to stress: when and how. J. Neuroendocrinol. 15: 711–724. https://doi.org/10.1046/j.1365-2826.2003.01033.x

Wingfield, J. C., Romero, L. M., 2001. Adrenocortical responses to stress and their modulation in free-living vertebrates. Handbook of physiology pp. 211–236. https://doi.org/10.1002/cphy.cp070411

Zbyryt, A., Bubnicki, J.W., Kuijper, D.P.J., Dehnhard, M., Churski, M., Schmidt, K. 2017. Do wild ungulates experience higher stress with humans than with large carnivores? Behav. Ecol. 29: 19–30. https://doi.org/10.1093/beheco/arx142

