## Supplementary material for "Age and spatio-temporal variations in food resources modulate stress-immunity relationships in three populations of wild roe deer": Model selection Chize

**Table S3**: Performance of the subset of candidate linear mixed-effect models within a ΔAICc < 2 fitted to investigate variation in immune parameters in the roe deer population of Chizé. Model(s) in bold was/were used for estimation of parameters and averaged when more than one model was considered after removing models that differed from a higher-ranking model by the addition of one or more parameters. These were rejected as uninformative, as recommended by Arnold (2010) and Richards (2008). Our set of candidate models was composed of all simpler models that included faecal cortisol metabolites (FCMs), sex, body weight, age, local resources quality and quantity (sector), temporal variations of resources among years (year quality), the time delay between capture and blood sampling (delay), Julian date of capture (Julian date), the two-way interaction between FCMs and age, the two-way interaction between FCMs and sector, and between FCMs and quality of the year, the two-way interaction between FCMs and weight, and between FCMs and sex, and finally the two-way interaction between sex and age. Individual identity, year of capture, and birth cohort were included as random effects. AICc is the value of the corrected Akaike’s Information Criterion and K is the number of estimated parameters for each model. The ranking of the models is based on the differences in the values for ΔAICc and on the Akaike weights (AICw).

A) Neutrophiles

| **Models** | **K** | **AICc** | **ΔAICc** | **AICw** |
| --- | --- | --- | --- | --- |
| **Age+Julian day+Delay+FCMs+Year quality** | **10** | **1574.1** | **0.00** | **0.055** |
| **Age+Julian day+Delay+FCMs** | **9** | **1574.2** | **0.13** | **0.051** |
| **Age^2+Julian day+Delay+FCMs+Weight+Year quality** | **11** | **1574.2** | **0.15** | **0.051** |
| Age^2+Julian day+Delay+FCMs+Weight+Year quality+Sex | 12 | 1574.4 | 0.34 | 0.046 |
| Age+Julian day+Delay+FCMs+Weight+Year quality | 11 | 1574.6 | 0.52 | 0.042 |
| Age+Julian day+Delay+FCMs+Year quality+Sex | 11 | 1574.7 | 0.65 | 0.040 |
| **Age^2+Julian day+Delay+FCMs+Weight** | **10** | **1574.9** | **0.78** | **0.037** |
| Age+Julian day+Delay+FCMs+Weight+Year quality+Sex | 12 | 1574.9 | 0.80 | 0.037 |
| Age+Julian day+Delay+FCMs+Sex | 10 | 1574.9 | 0.85 | 0.036 |
| Age+Julian day+Delay+FCMs+Weight | 10 | 1575.0 | 0.90 | 0.035 |
| Age^2+Julian day+Delay+FCMs+Weight+Sex | 11 | 1575.1 | 1.04 | 0.033 |
| **Age+Julian day+Delay** | **8** | **1575.2** | **1.09** | **0.032** |
| **Age^2+Julian day+Delay+Weight+Year quality** | **10** | **1575.3** | **1.20** | **0.030** |
| Age+Julian day+Delay+FCMs+Weight+Sex | 11 | 1575.4 | 1.30 | 0.029 |
| Age^2+Julian day+Delay+Weight+Year quality+Sex | 11 | 1575.4 | 1.30 | 0.029 |
| **Age^2+Julian day+Delay+Weight** | **9** | **1575.5** | **1.40** | **0.027** |
| Age+Julian day+Delay+Year quality | 9 | 1575.5 | 1.41 | 0.027 |
| **Age^2+Delay+FCMs+Weight+Year quality** | **10** | **1575.6** | **1.48** | **0.026** |
| Age+Julian day+Delay+Weight | 9 | 1575.7 | 1.59 | 0.025 |
| Age^2+Julian day+Delay+Weight+Sex | 10 | 1575.7 | 1.61 | 0.025 |
| Age+Julian day+Delay+FCMs+Year quality+FCMs*Year quality | 11 | 1575.7 | 1.61 | 0.025 |
| Age+Julian Date+Delay+Weight+Year quality | 10 | 1575.7 | 1.61 | 0.024 |
| Age^2+Delay+FCMs+Weight+Year quality+Sex | 11 | 1575.7 | 1.64 | 0.024 |
| Age^2+Julian date+Delay+FCMs+Weight+Year quality+FCMs*Year quality | 12 | 1575.8 | 1.73 | 0.023 |
| Age+Julian date+Delay+Sex | 9 | 1575.8 | 1.76 | 0.023 |
| Age+Julian date+Delay+Weight+Year quality+Sex | 11 | 1575.9 | 1.79 | 0.022 |
| Age^2+Julian date+Delay+FCMs+Weight+Year quality+Sex+FCMs*Year quality | 13 | 1575.9 | 1.80 | 0.022 |
| Age^2+Julian date+Delay+FCMs+Weight+Year quality+FCMs*Year quality | 12 | 1575.9 | 1.85 | 0.022 |
| Age+Julian date+Delay+Weight+Sex | 10 | 1576.0 | 1.91 | 0.021 |
| Age+Julian date+Delay+Year quality+Sex | 10 | 1576.1 | 1.98 | 0.020 |
| **Age^2+Delay+Weight+Year quality** | **9** | **1576.1** | **1.98** | **0.020** |
| Age+Age^2+Julian date+Delay+FCMs+Year quality | 11 | 1576.1 | 1.99 | 0.020 |
| Age^2+Julian date+Delay+FCMs+Weight+Year quality+Sex+FCMs*Sex | 13 | 1576.1 | 2.00 | 0.020 |

B) Eosinophiles

| **Models** | **K** | **AICc** | **ΔAICc** | **AICw** |
| --- | --- | --- | --- | --- |
| **Age^2+Delay+Weight+Year quality+Sector+Sex+Age^2*Sex** | **13** | **-601.6** | **0.00** | **0.238** |
| **Age+Age^2+Delay+Weight+Year quality+Sector** | **12** | **-600.7** | **0.91** | **0.151** |
| Age^2+Julian day+Delay+Weight+Year quality+Sector+Sex+Age^2*Sex | 14 | -600.6 | 1.03 | 0.142 |
| **Age^2+Delay+Weight+Year quality+Sector** | **11** | **-605.5** | **1.13** | **0.135** |
| Age+Age^2+Delay+Weight+Year quality+Sector+Sex+Age^2*Sex | 14 | -600.3 | 1.32 | 0.123 |
| **Delay+Weight+Year quality+Sector** | **10** | **-600.1** | **1.50** | **0.112** |
| Age^2+Delay+Weight+Year quality+Sector+Sex+Age^2*Sex+FCMs | 14 | -599.8 | 1.7 | 0.099 |

C) Basophiles

| **Models** | **K** | **AICc** | **ΔAICc** | **AICw** |
| --- | --- | --- | --- | --- |
| **FCMs+Weight+Sector+FCMs*Sector** | **11** | **-1065.8** | **0.00** | **0.077** |
| **Weight** | **6** | **-1065.8** | **0.01** | **0.077** |
| **Age+FCMs+Sector+FCMs*Sector** | **11** | **-1065.7** | **0.17** | **0.071** |
| FCMs+Weight | 7 | -1065.4 | 0.47 | 0.061 |
| Age+Age^2+FCMs+Sector+FCMs*Sector | 12 | -1065.2 | 0.64 | 0.056 |
| Age+FCMs+Weight+Sector+FCMs*Sector | 12 | -1064.9 | 0.88 | 0.050 |
| **Age** | **6** | **-1064.7** | **1.11** | **0.044** |
| Age+FCMs | 7 | -1064.6 | 1.22 | 0.042 |
| Age^2+FCMs+Weight+Sector+FCMs*Sector | 12 | -1064.5 | 1.33 | 0.040 |
| Age+Weight | 7 | -1064.4 | 1.47 | 0.037 |
| Age+Age^2 | 7 | -1064.2 | 1.59 | 0.035 |
| Age+FCMs+Sector+Sex+FCMs*Sector | 12 | -1064.2 | 1.64 | 0.034 |
| Age+Age^2+FCMs | 8 | -1064.2 | 1.66 | 0.034 |
| Julian date+FCMs+Weight+Sector+FCMs*Sector | 12 | -1064.1 | 1.68 | 0.033 |
| Age+Julian date+FCMs+Sector+FCMs*Sector | 12 | -1064.1 | 1.69 | 0.033 |
| Age^2+Weight | 7 | -1064.1 | 1.72 | 0.033 |
| Julian date+Weight | 7 | -1064.1 | 1.73 | 0.033 |
| Age+FCMs+Weight | 8 | -1064.0 | 1.82 | 0.031 |
| Weight+Sex | 7 | -1064.0 | 1.83 | 0.031 |
| FCMs+Weight+Sector+FCMs*Weight+FCMs*Sector | 12 | -1064.0 | 1.84 | 0.031 |
| Weight+Year quality | 7 | -1064.0 | 1.85 | 0.031 |
| Delay+FCMs+Weight+Sector+FCMs*Sector | 12 | -1063.9 | 1.91 | 0.030 |
| FCMs+Weight+Year quality+Sector+FCMs*Sector | 12 | -1063.9 | 1.95 | 0.029 |
| FCMs+Weight+Sector+Sex+FCMs*Sector | 12 | -1063.8 | 2.00 | 0.028 |

D) Monocytes

| **Models** | **K** | **AICc** | **ΔAICc** | **AICw** |
| --- | --- | --- | --- | --- |
| **Age+FCMs+Year quality+Sex+FCMs*Year quality** | **9** | **-187.4** | **0.00** | **0.276** |
| Age+Julian date+FCMs+Year quality+Sex+FCMs*Year quality | 10 | -186.2 | 1.24 | 0.149 |
| Age+Delay+FCMs+Year quality+Sex+FCMs*Year quality | 10 | -185.9 | 1.52 | 0.129 |
| Age+FCMs+Weight+Year quality+Sex+FCMs*Year quality | 10 | -185.8 | 1.61 | 0.123 |
| Age+Age^2+FCMs+Year quality+Sex+FCMs*Year quality | 10 | -185.7 | 1.74 | 0.116 |
| Age+FCMs+Year quality+Sex+Age*FCMs+FCMs*Year quality | 10 | -185.5 | 1.94 | 0.105 |
| **Age+FCMs+Year quality+Sex** | **8** | **-185.4** | **2.00** | **0.102** |

E) Hemagglutination

| **Models** | **K** | **AICc** | **ΔAICc** | **AICw** |
| --- | --- | --- | --- | --- |
| **Age+Julian date+ FCMs+Weight+age*FCMs** | **10** | **1057.8** | **0.00** | **0.407** |
| **Age+ FCMs+Weight+age*FCMs** | **9** | **1058.7** | **0.84** | **0.268** |
| Age+Julian date+ FCMs+Weight+Year quality+age*FCMs | 11 | 1059.6 | 1.81 | 0.164 |
| Age+Julian date+ FCMs+Weight+age*FCMs+FCMs*Weight | 11 | 1059.7 | 1.86 | 0.161 |

F) Hemolysis

| **Models** | **K** | **AICc** | **ΔAICc** | **AICw** |
| --- | --- | --- | --- | --- |
| **Age+Julian date** | **7** | **868.7** | **0.00** | **0.217** |
| Age+Julian date+Delay | 8 | 869.4 | 0.66 | 0.156 |
| Age+Julian date+Year quality | 8 | 869.5 | 0.79 | 0.146 |
| **Julian date** | **6** | **869.9** | **1.20** | **0.119** |
| Age+Julian date+Delay+Year quality | 9 | 870.2 | 1.51 | 0.102 |
| Age+Julian date+FCMs+Age*FCMs | 9 | 870.4 | 1.71 | 0.092 |
| Julian date+Delay | **7** | 870.5 | 1.85 | 0.086 |
| Age+Julian date+Weight | 8 | 870.7 | 1.98 | 0.081 |

G) Gamma globulins

| **Models** | **K** | **AICc** | **ΔAICc** | **AICw** |
| --- | --- | --- | --- | --- |
| **Age+Delay+FCMs+Sector+age*FCMs+Sector*FCMs** | **13** | **2334.0** | **0.00** | **0.441** |
| **Age+FCMs+Sector+age*FCMs+Sector*FCMs** | **12** | **2335.4** | **1.41** | **0.218** |
| Age+Delay+FCMs+Sector+Sex+age*FCMs+Sector*FCMs | 14 | 2335.9 | 1.87 | 0.173 |
| Age+Delay+FCMs+Year quality+Sector+age*FCMs+Sector*FCMs | 14 | 2335.9 | 1.93 | 0.168 |

H) Lymphocytes

| **Models** | **K** | **AICc** | **ΔAICc** | **AICw** |
| --- | --- | --- | --- | --- |
| **Age+Age^2+Delay+FCMs+Year quality+Sex+I(Age^2)*Sex+FCMs*Year quality** | **13** | **41.3** | **0.00** | **0.344** |
| **Age+Age^2+Delay+Sex+I(Age^2)*Sex+FCMs*Year quality** | **10** | **41.3** | **0.00** | **0.344** |
| Age+Age^2+Delay+FCMs+Sex+I(Age^2)*Sex | 11 | 42.8 | 1.56 | 0.158 |
| Age+Age^2+Delay+FCMs+Year quality+Sex+I(Age^2)*Sex+FCMs*Year quality | 14 | 42.9 | 1.61 | 0.154 |

I) Alpha-1 globulins

| **Models** | **K** | **AICc** | **ΔAICc** | **AICw** |
| --- | --- | --- | --- | --- |
| **Delay+FCMs+Weight+Sex** | **9** | **-567.5** | **0.00** | **0.082** |
| Delay+FCMs+Weight+Sex+FCMs*Sex | 10 | -567.2 | 0.25 | 0.072 |
| Delay+FCMs+Weight+Year quality+Sex+FCMs*Year quality | 11 | -567.1 | 0.37 | 0.068 |
| Delay+FCMs+Weight+Year quality+Sex+FCMs*Year quality+FCMs*Sex+FCMs*Year quality | 12 | -566.9 | 0.56 | 0.062 |
| Age^2+Delay+FCMs+Weight+Sex+I(Age^2)*Sex | 11 | -566.6 | 0.86 | 0.053 |
| Age+Age^2+Delay+FCMs+Weight+Sex+I(Age^2)*Sex | 12 | -566.2 | 1.27 | 0.043 |
| Age^2+Delay+FCMs+Weight+Year quality+Sex+I(Age^2)*Sex+FCMs*Year quality | 13 | -566.2 | 1.28 | 0.043 |
| Delay+FCMs+Weight+Year quality+Sex | 10 | -566.2 | 1.33 | 0.042 |
| Age+Age^2+Delay+FCMs+Weight+Year quality+Sex+Age*FCMs+I(Age^2)*Sex+FCMs*Year quality | 15 | -566.1 | 1.33 | 0.042 |
| Age^2+Delay+FCMs+Weight+Sector+Sex+I(Age^2)*Sex+FCMs*Sector | 15 | -566.1 | 1.37 | 0.041 |
| Delay+FCMs+Weight+Sector+Sex+FCMs*Sector | 13 | -566.1 | 1.40 | 0.041 |
| Age+Age^2+Delay+FCMs+Weight+Year quality+Sex+I(Age^2)*Sex+FCMs*Year quality | 14 | -566.1 | 1.42 | 0.040 |
| Age+Age^2+Delay+FCMs+Weight+Sector+Sex+I(Age^2)*Sex+FCMs*Sector | 16 | -566.0 | 1.52 | 0.038 |
| Age+Age^2+Delay+FCMs+Weight+Year quality+Sector+Sex+I(Age^2)*Sex+FCMs*Year quality+FCMs*Sector | 18 | -565.9 | 1.61 | 0.037 |
| Delay+FCMs+Weight+Year quality+Sex+FCMs*Sex | 11 | -565.8 | 1.63 | 0.036 |
| Age+Age^2+Delay+FCMs+Weight+Sex+Age*FCMs+I(Age^2)*Sex | 13 | -565.8 | 1.66 | 0.036 |
| Age^2+Delay+FCMs+Weight+Sex+I(Age^2)*Sex+FCMs*Sex | 12 | -565.8 | 1.69 | 0.035 |
| Age^2+Delay+FCMs+Weight+Year quality+Sector+Sex+I(Age^2)*Sex+FCMs*Year quality+FCMs*Sector | 17 | -565.7 | 1.81 | 0.033 |
| Delay+FCMs+Weight+Year quality+Sector+Sex+FCMs*Year quality+FCMs*Sector | 15 | -565.7 | 1.81 | 0.033 |
| Julian date+Delay+FCMs+Weight+Sex | 10 | -565.6 | 1.93 | 0.031 |
| Age+Age^2+Delay+FCMs+Weight+Year quality+Sector+Sex+Age*FCMs+I(Age^2)*Sex+FCMs*Year quality+FCMs*Sector | 19 | -565.5 | 1.96 | 0.031 |
| Age^2+Delay+FCMs+Weight+Sex | 10 | -565.5 | 1.97 | 0.031 |
| Age+Age^2+Delay+FCMs+Weight+Sex+I(Age^2)*Sex+FCMs*Sex | 13 | -565.5 | 1.98 | 0.030 |

J) Alpha-2 globulins

| **Models** | **K** | **AICc** | **ΔAICc** | **AICw** |
| --- | --- | --- | --- | --- |
| **Julian date+Delay+FCMs+Weight+Year quality+Sector+FCMs*Year quality+FCMs*Sector** | **15** | **-282.3** | **0.00** | **0.222** |
| **Julian date+FCMs+Weight+Year quality+Sector+FCMs*Year quality+FCMs*Sector** | **14** | **-281.9** | **0.47** | **0.176** |
| **Julian date+Delay+FCMs+Year quality+Sector+FCMs*Year quality+FCMs*Sector** | **14** | **-281.5** | **0.84** | **0.146** |
| Julian date+Delay+FCMs+Weight+Year quality+Sector+FCMS*Weight+FCMs*Year quality+FCMs*Sector | 16 | -280.7 | 1.61 | 0.099 |
| **Julian date+FCMs+Year quality+Sector+FCMs*Year quality+FCMs*Sector** | **13** | **-280.7** | **1.62** | **0.099** |
| **Delay+FCMs+Weight+Year quality+Sector+FCMs*Year quality+FCMs*Sector** | **14** | **-280.5** | **1.82** | **0.090** |
| Julian date+Delay+FCMs+Weight+Year quality+Sector+Sex+FCMs*Year quality+FCMs*Sector | 16 | -280.4 | 1.94 | 0.084 |
| **FCMs+Weight+Year quality+Sector+FCMs*Year quality+FCMs*Sector** | **13** | **-280.4** | **1.96** | **0.083** |

K) Beta globulins

| **Models** | **K** | **AICc** | **ΔAICc** | **AICw** |
| --- | --- | --- | --- | --- |
| **Age+Age^2+Julian date+FCMs+Weight+Sector+Sex** | **15** | **-368.9** | **0.00** | **0.289** |
| Age+Age^2+Julian date+FCMs+Weight+Sector+Sex+Age*FCMs | 16 | -368.1 | 0.84 | 0.190 |
| Age+Age^2+Julian date+FCMs+Weight+Sector+Sex+I(Age^2)*Sex | 16 | -367.6 | 1.27 | 0.153 |
| Age+Age^2+Julian date+Delay+FCMs+Weight+Sector+Sex | 16 | -367.3 | 1.59 | 0.130 |
| **Age+Age^2+Julian date+Weight+Sex** | **10** | **-367.2** | **1.74** | **0.121** |
| Age+Age^2+Julian date+FCMs+Weight+Year quality+Sector+Sex | 16 | -367.1 | 1.81 | 0.117 |

L) Haptoglobin

| **Models** | **K** | **AICc** | **ΔAICc** | **AICw** |
| --- | --- | --- | --- | --- |
| **Julian Date+Year quality+Sex** | **8** | **297.6** | **0.00** | **0.190** |
| Julian Date+Delay+Year quality+Sex | 9 | 298.0 | 0.45 | 0.152 |
| Julian Date+FCMs+Year quality+Sex+FCMs*Year quality | 10 | 298.4 | 0.85 | 0.124 |
| Age+Julian Date+Year quality+Sex | 9 | 298.7 | 1.12 | 0.109 |
| Julian date+Delay+FCMs+Year quality+Sex+FCMs*Year quality | 11 | 298.9 | 1.30 | 0.099 |
| Julian date+FCMs+Year quality+Sex | 9 | 299.1 | 1.50 | 0.090 |
| Age+Julian date+Delay+Year quality+Sex | 10 | 299.2 | 1.63 | 0.084 |
| Julian date+FCMs+Year quality+sex+FCMs*Year quality+FCMs*Sex | 11 | 299.4 | 1.83 | 0.076 |
| Date julian+Weight+Year quality+Sex+ | 9 | 299.4 | 1.88 | 0.075 |
