## Supplementary material for "Age and spatio-temporal variations in food resources modulate stress-immunity relationships in three populations of wild roe deer": Correlation matrix

**Table S5**: Correlation matrix for each population (Aurignac, Chizé, Trois-Fontaines) between the twelve immune parameters considered in our study.

A) Aurignac

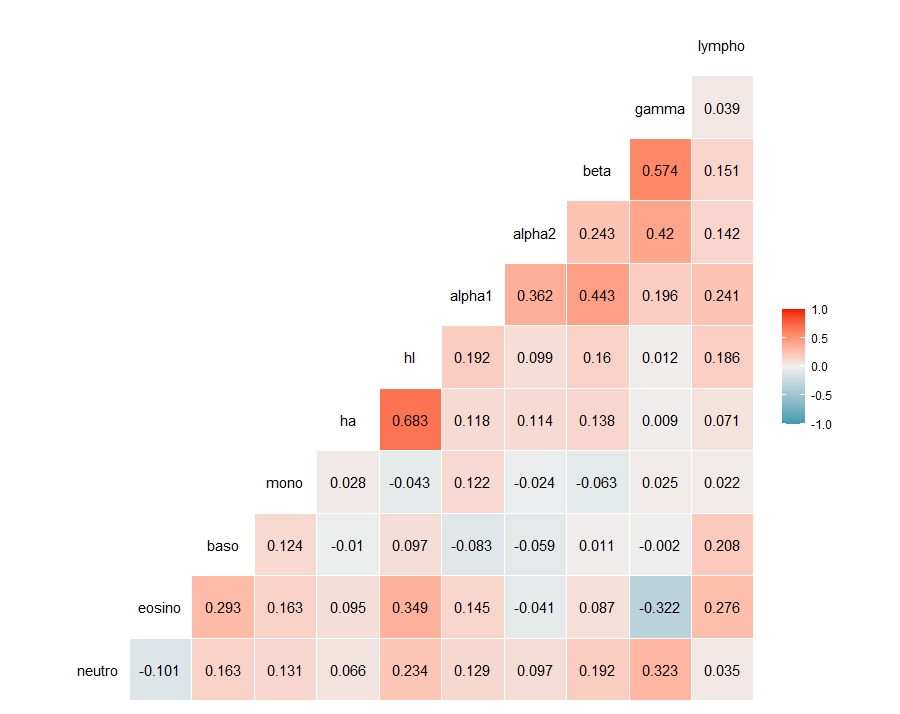

B) Chizé

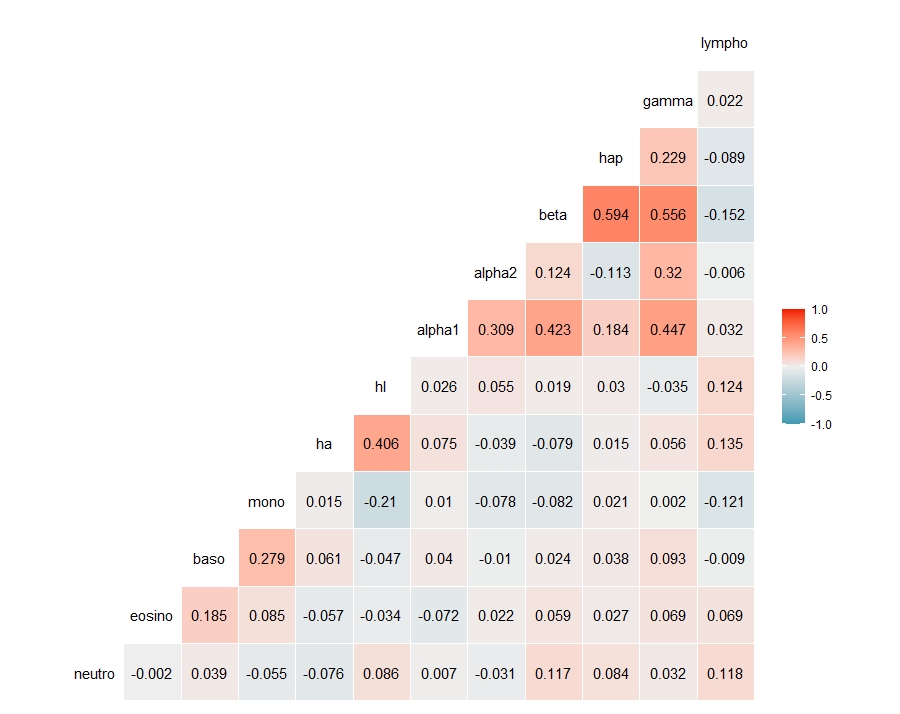

C) Trois-Fontaines

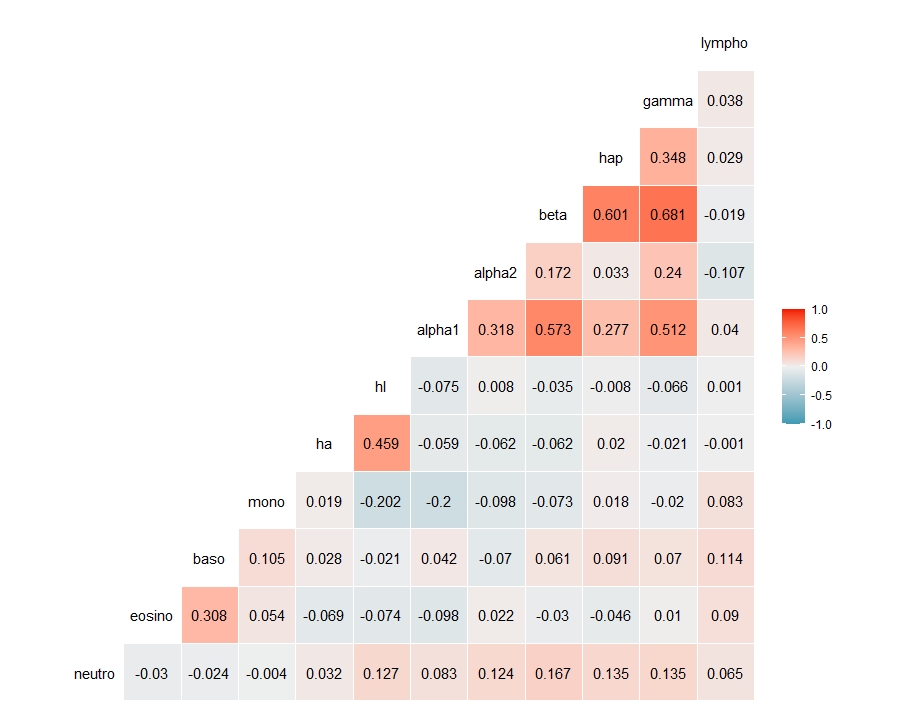
