## Supplementary material for "Age and spatio-temporal variations in food resources modulate stress-immunity relationships in three populations of wild roe deer": Predictions

|  | | **A** | **C** | **TF** | **A** | **C** | **TF** | **A** | **C** | **TF** | **A** | **C** | **TF** |
| --- | --- | --- | --- | --- | --- | --- | --- | --- | --- | --- | --- | --- | --- |
|  |  | (i) FCMs | | | (ii) Sector | | | (ii) Year | | | (iii) Age | | |
| Innate cellular | Neutrophil | -0.51 | -0.26 | 0 | 0 | 0 | NA | 0 | 0 | 0 | 0 | 0 | 0 |
|  | Eosinophil | 0 | 0 | 0 | 0 | 0 |  | 0 | 0 | 0 | 0 | 0 | 0 |
|  | Basophil | 0 | -0.002 | 0 | 0 | -0.03 |  | 0 | 0 | 0 | 0 | 0 | 0 |
|  | Monocytes | 0 | -0.04 | -0.06 | 0 | 0 |  | 0 | 0.06 | 0 | 0 | 0 | 0 |
| Innate humoral | Hemagglutination | 0 | 2.57 | 0 | 0 | 0 |  | 0.96 | 0 | 0 | 0 | -0.3 | 0 |
|  | Hemolysis | -0.19 | 0 | 0 | 0 | 0 |  | 0 | 0 | 0 | 0 | 0 | 0 |
| Inflammatory markers | Alpha-1 globulin | 0 | -0.03 | 0 | -0.03 | 0 |  | 0 | 0 | 0 | 0 | 0 | 0 |
|  | Alpha-2 globulin | 0 | -0.004 | 0 | 0 | -0.06 |  | 0 | -0.06 | 0 | 0 | 0 | 0 |
|  | Beta globulin | 0.03 | 0 | 0 | 0 | 0.11 |  | 0 | 0 | 0 | 0 | 0 | 0 |
|  | Haptoglobin | NA | 0 | 0 | NA | 0 |  | NA | 0 | 0 | NA | 0 | -0.08 |
| Adaptive cellular | Lymphocytes | 0 | 0 | 0 | 0 | 0 |  | 0 | -0.09 | 0 | 0 | 0 | -0.03 |
| Adaptive humoral | Gamma globulin | 1.58 | 1.13 | 0 | -2.11 | 1.77 |  | 0 | 0 | 0 | 0 | -0.30 | 0 |

**Table S4 –** Summary of the results from our study in relation to our predictions for each immune parameter considered in the three studied populations: Aurignac (A), Chizé (C), and Trois-Fontaines (TF). Results that are in accordance with our predictions are indicated in green, those that are opposite to our predictions are indicated in red, and no colour means that there was no relationship between the considered immune parameter and either FCMs (prediction i), the two-way interaction between FCMs and sector identity (prediction ii), the two-way interaction between FCMs and year quality (prediction ii), or the two-way interaction between FCMs and age (prediction iii). In the case of significant effects of the two-way interaction between FCMs and sector identity, the estimate’s values reported are the one of the lowest quality sector (sector 3). Sector identity indexes spatial heterogeneity in food resources, while year quality indexes temporal heterogeneity in food resources.
